# EARLY FLOWERING 3 (ELF3): a novel role in integrating environmental stimuli with root stem cell niche maintenance

**DOI:** 10.64898/2026.01.12.699078

**Authors:** Ali Eljebbawi, Rebecca C. Burkart, Laura Czempik, Vivien I. Strotmann, Lorelei Masselot-Joubert, Audrey Guillotin, Xuelei Lai, Mark d. Tully, Chloe Zubieta, Stephanie Hutin, Yvonne Stahl

## Abstract

Maintaining stem cell niche (SCN) homeostasis in the root apical meristem (RAM) is essential for proper root growth and thus for overall plant development. In *Arabidopsis thaliana*, a group of slowly dividing cells at the SCN center known as the quiescent center (QC) maintain the surrounding stem cells, including the distally located columella stem cells (CSCs) which give rise to the differentiated columella cells. Many actors, including the PLETHORA (PLT) family transcription factors, regulate the QC quiescence and CSC fate. However, little is known about the integration of external and/or internal cues into regulating SCN homeostasis. In this study, we report for the first time the interaction between PLT3 and the thermosensor and circadian clock related transcriptional regulator, EARLY FLOWERING 3 (ELF3), which are both expressed in the root SCN. We show that ELF3 localizes, similar to PLT3, to subcellular condensates and sustains the QC and CSC fate. We demonstrate that ELF3 forms condensates *in vitro* and *in vivo*, in the cytoplasm, as well as in the nucleus, where it then co-localizes with PLT3. We reveal that the interaction of ELF3 and PLT3 is mediated by their intrinsically disordered prion-like domains (PrDs). Furthermore, transient expression in human epithelial cells (HEp-2) cells and in *Nicotiana benthamiana* shows that PHYTOCHROME INTERACTING FACTORS 3 and 4 (PIF3/4) function as nuclear shuttles for ELF3, recruiting it to nuclear condensates, where it co-localizes with PLT3, PIF3, and PIF4. Accordingly, we propose a model where the co-localization and interactions of ELF3, PLT3, PIF3, and PIF4 represent a dynamic mechanism to integrate environmental signals into SCN maintenance and cell fate decisions.

## Introduction

Plants require well-developed and functional root systems for anchorage in the soil and the uptake of water and nutrients. In *Arabidopsis thaliana* (*A. thaliana)*, root development is supported by a group of pluripotent stem cells forming the stem cell niche (SCN) in the root apical meristem (RAM) at the root apex. Structurally, the root SCN consists of a central cluster of four to eight rarely dividing cells forming the quiescent center (QC), surrounded by a layer of faster dividing stem cell initials (Dolan et al. 1993; Lu et al. 2021) (Figure 1a). These initials are organized into distinct populations including the columella stem cells (CSCs) distal to the QC, the cortex/endodermis initials, the epidermis/lateral root cap initials, and the stele initials that give rise to the vascular tissues and pericycle. Functionally, the QC serves as a long-term stem cell reservoir and prevents the premature differentiation of the surrounding initials. In turn, the initials divide and differentiate to produce the necessary and distinct root tissues. For instance, the distally located CSCs differentiate into columella cells (CCs), which accumulate starch granules, known as statoliths, and specialize in gravity sensing. Maintaining the SCN homeostasis requires a tight balance between cell division and differentiation, which is intricately regulated by phytohormones and transcription factors (TFs) (Benfey et al. 1993; van den Berg et al. 1995; van den Berg et al. 1997; Drisch and Stahl 2015; Yazdanbakhsh et al. 2011; Fisher and Sozzani 2016). One key regulator of SCN maintenance is the homeodomain TF WUSCHEL-RELATED-HOMEOBOX5 (WOX5). WOX5 is expressed in the QC and maintains the surrounding initials via non-cell-autonomous signaling (Sarkar et al. 2007; Pi et al. 2015). Additionally, the APETALA2-type TFs of the PLETHORA family (PLTs) are also known for their role in SCN maintenance. Four *PLTs* (*PLT1*, *PLT2*, *PLT3*, and *PLT4*) are expressed in the RAM, with their expression levels are tightly correlated to local auxin concentrations. Accordingly, they form a concentration-gradient which peaks around the QC maintaining its quiescence. This is evident in multiple *plt* mutants which show defective SCN phenotypes with increased QC divisions and stem cell differentiation, as shown in *plt1*, *plt2* and *plt2*, *plt3* (Aida et al. 2004; Galinha et al. 2007; Burkart et al. 2022). Recently, we have shown that the PLT3 and WOX5 interaction maintains the distal SCN in an interdependent way, as they form protein complexes within subnuclear condensates (Burkart et al. 2022).

**Figure 1:**
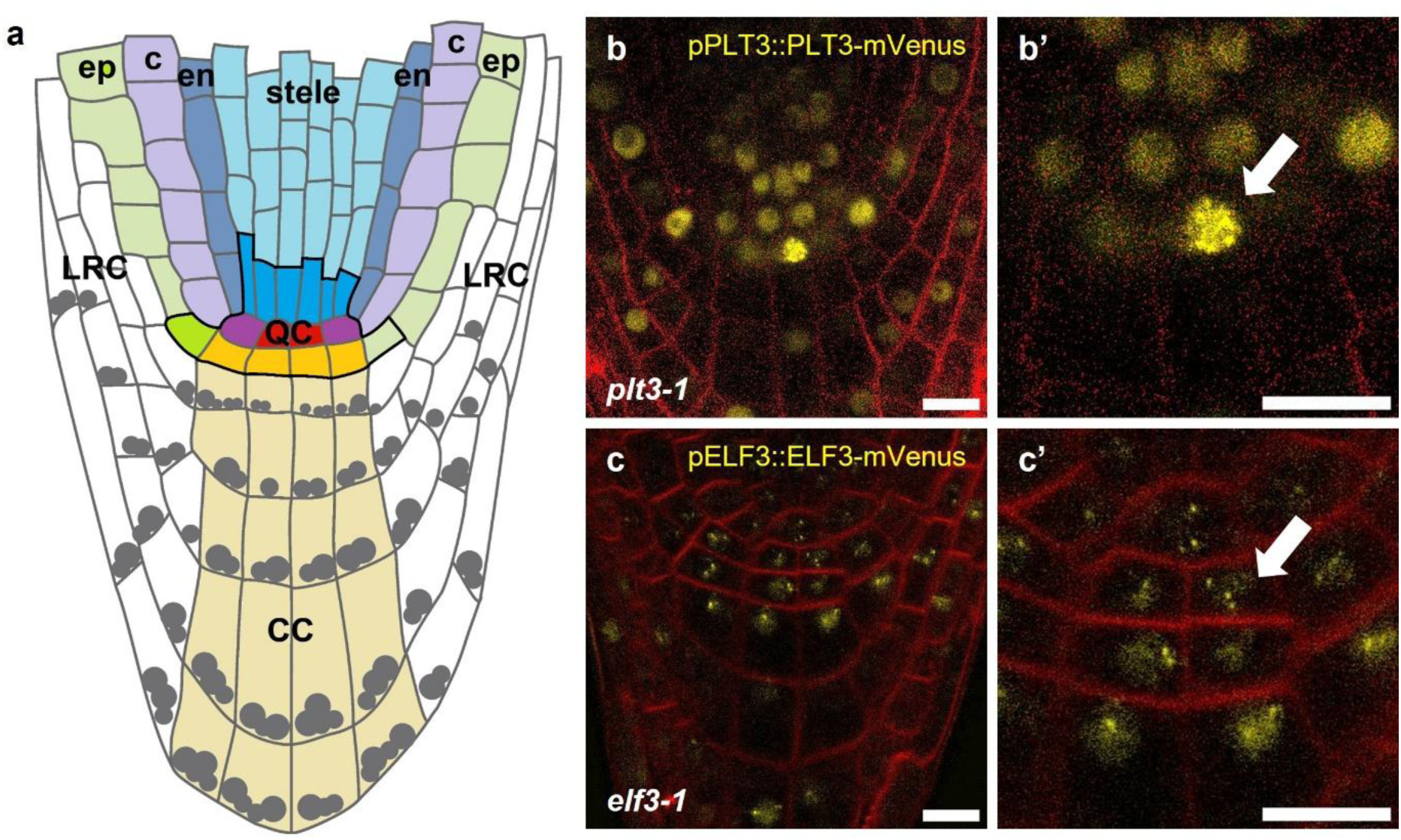
Localization of PLT3 and ELF3 in the SCN of *Arabidopsis thaliana*. **a:** Schematic representation of the *Arabidopsis thaliana* root apical meristem (RAM). The QC cells (red) maintain the surrounding stem cells (initials), together building the root stem cell niche (SCN) outlined in black. The different cell types are color-coded. QC= quiescent centre (red); CC= columella cells (light orange); LRC= lateral root cap (white); ep= epidermis (light green); c= cortex (light purple); en= endodermis (dark blue); cortex/endodermis initials (purple); grey dots= starch granules. **b, c:** Representative images of the SCN of *A. thaliana* seedlings in *plt3-1* or *elf3-1* background respectively expressing pPLT3::PLT3-mVenus (b) and pELF3::ELF3-mVenus (c) (yellow) with FM4-64 staining (red). **b’, c’:** Magnifications of **b, c** with arrowheads pointing at condensates. Scale bars represent 10 µm.

While additional factors involved in this process remain to be discovered, a key open question is how internal cues, like circadian rhythms, and external cues, such as light and temperature, are integrated into root growth and SCN maintenance. The circadian clock is a key developmental regulator that synchronizes growth with daily and seasonal changes not only in aerial tissues but also in roots (Yazdanbakhsh et al. 2011; Sanchez and Kay 2016). For instance, oscillations in root elongation have been linked to the circadian control of carbon partitioning (Yazdanbakhsh et al. 2011). Temperature is also an essential factor that affects root development by regulating cell division and elongation in the root meristem, shaping root growth and architecture (Pyl et al. 2012; Yang et al. 2017). The temperature sensing transcriptional regulator EARLY FLOWERING 3 (ELF3), a core component of the circadian clock, is expressed not only in aerial tissues, but also in the root (Jung et al. 2020). ELF3 controls the rhythmic development of roots and links the shoot and root responses to environmental stimuli (James et al. 2008; Yazdanbakhsh et al. 2011; Joseph et al. 2015; Li et al. 2019). Initially identified for its role in regulating flowering time, as *elf3* knockout mutants display early-flowering phenotypes and insensitivity to photoperiods (Zagotta et al. 1996; Hicks 2001), recent studies have also shown that ELF3 plays an important role in temperature sensing and response (Jung et al. 2020). ELF3, LUX ARRYTHMO, and ELF4 form the temperature sensitive transcriptional repressor evening complex (EC) and regulate expression of target genes according to circadian rhythms (Covington 2001; Hicks 2001; Liu 2001; Silva et al. 2020). The EC plays a role in thermomorphogenesis by suppressing the transcription of *PHYTOCHROME INTERACTING FACTOR 4* (*PIF4*) and *PHYTOCHROME INTERACTING FACTOR 5* (*PIF5*) (Nusinow et al. 2011), in a temperature-dependent manner. The PIFs are basic helix-loop-helix (bHLH) TFs that modulate the expression of light-responsive genes and transmit light signals that the phytochromes (PHYs) and cryptochromes (CRYs) receive (Pedmale et al. 2016; Cordeiro et al. 2022). They play key roles in the shade avoidance response and thermomorphogenesis by directly regulating genes important for elongation growth and hormone signaling (Koini et al. 2009; Zhao and Bao 2021). Remarkably, PIFs colocalize with several regulators, including PHYs, CRYs, and ELF3, in subcellular condensates known as photobodies (Franklin et al. 2011). Moreover, PIFs interact with FLOWERING CONTROL LOCUS A (FCA), AUXIN RESPONSE FACTOR 6 (ARF6), and many other regulators. These interactions integrate diverse environmental stimuli and regulate several developmental processes (Leivar et al. 2008a; Leivar et al. 2008b; Qiu et al. 2017; Jiang et al. 2019).

Interestingly, the ELF3 and PLT3 proteins share common features, where both contain intrinsically disordered regions (IDRs) including prion-like domains (PrD) that are enriched in glutamines, methionines, and tyrosines (Burkart et al. 2022; Hutin et al. 2023). PrDs can mediate multivalent interactions, protein-protein interactions and drive phase separation, contributing to compartmentalization and several cellular processes in plant cells (Eljebbawi et al. 2023; Legen et al. 2024; Wu and Li 2024; Peng et al. 2025; Hutin et al. 2026). ELF3 serves as a scaffold that binds a variety of transcription factors (TFs), and it undergoes condensation driven by its PrD (Hutin et al. 2023). ELF3 condensation is reversible and is triggered by elevated temperatures as shown in *A. thaliana* and *in vitro* experiments (Jung et al. 2020; Hutin et al. 2023). Our previous results have demonstrated that PLT3 also forms condensates with the TF, WOX5 (Burkart et al. 2022). Due to their overlapping expression patterns, the presence of a PrD, the shared property of condensate formation and ELF3’s role as a promiscuous scaffold protein, we investigated the interactions of ELF3 with PLT3 as well as PIF3 and PIF4. We showed that it directly interacts with all three proteins and forms nuclear condensates. Based on these results, we propose a model in which ELF3 acts as an integrator of light and temperature signals into root growth, where it plays a role along with PLT3 in SCN maintenance.

## Results

### Both ELF3 and PLT3 are expressed in the RAM of *Arabidopsis thaliana*

We analyzed the expression patterns of *PLT3* and *ELF3* in the *A. thaliana* root SCN (Figure 1a) using fluorescent protein (FP)-tagged fusion proteins driven by their native promoters. In the *pPLT3::PLT3*-*mVenus* reporter line in *plt3-1* mutant background (Burkart et al. 2022), a concentration gradient of PLT3 was observed, with the highest concentration in the QC and CSC layer. As previously described, PLT3 nuclear condensates were detected in the CSC layer (Figure 1b-b’) (Burkart et al 2022). To determine ELF3 localization in the root, we produced a stable *pELF3::ELF3-mVenus* reporter line in *elf3-1* mutant plants and found that ELF3 is ubiquitously expressed in all RAM cells, including the SCN. Notably, ELF3 accumulates in nuclear condensates within all SCN root cells (Figure 1c-c’). The overlapping expression domains of PLT3 and ELF3 in the SCN, as well as their mutual tendency to form condensates, prompted us to investigate their potential interconnection and putative roles in SCN maintenance. Therefore, the QC and CSC phenotypes in the respective mutants were analyzed. Importantly, the functionality of the fusion proteins was validated by their ability to complement the previously described phenotypes of the *elf3-1* mutant, including the previously described elongated hypocotyls and altered leaf development (Supplementary figure 1, Supplementary tables 11-13) (Zagotta et al. 1992; Hicks et al. 1996; Zagotta et al. 1996; Hicks 2001).

### *elf3* and *plt3* single and double mutants show defects in the *A. thaliana* root SCN

Next, we investigated whether ELF3 influences SCN homeostasis, potentially together or in parallel with PLT3. Therefore, the root meristem phenotypes of *A. thaliana* Col-0 wildtype seedlings, as well *plt3* and *elf3* single and *plt3 elf3* double mutants were analyzed. For this, we implemented SCN staining to simultaneously visualize cell divisions, starch granules, and cell walls within the same root, as previously described (Burkart et al. 2022).

Our experimental observations on the SCN of six day old seedlings showed that the CSC phenotypes were only mildly different between genotypes. In Col-0, the majority of the measured roots (67 %) had one CSC layer, and only 20 % displayed higher differentiation with zero CSC layers, consistent with previous findings (Burkart et al 2022, Strotmann et al 2025). As for *elf3-1* and *plt3-1* single mutants, 52 % and 56 % of their roots had one CSC layer, respectively. Both mutants showed a tendency towards increased differentiation, with 27-31 % of their roots lacking a CSC layer, 1.5-fold more than in Col-0. Notably, *elf3-1* roots additionally exhibited a distinct phenotype at the other end of the differentiation spectrum, by the presence of two CSC layers (21 %), compared to Col-0 and *plt3-1* (both 12 %), indicating reduced differentiation in this subset of roots. The *elf3-1 plt3-1* double mutant was similar to the single mutants, with 49 % of its roots having one CSC layer (Figure 2a-f). Nevertheless, the Kruskal-Wallis test was not significant (p-value = 0.236), suggesting that ELF3 and PLT3 do not substantially or additively change the average number of CSC layers.

**Figure 2:**
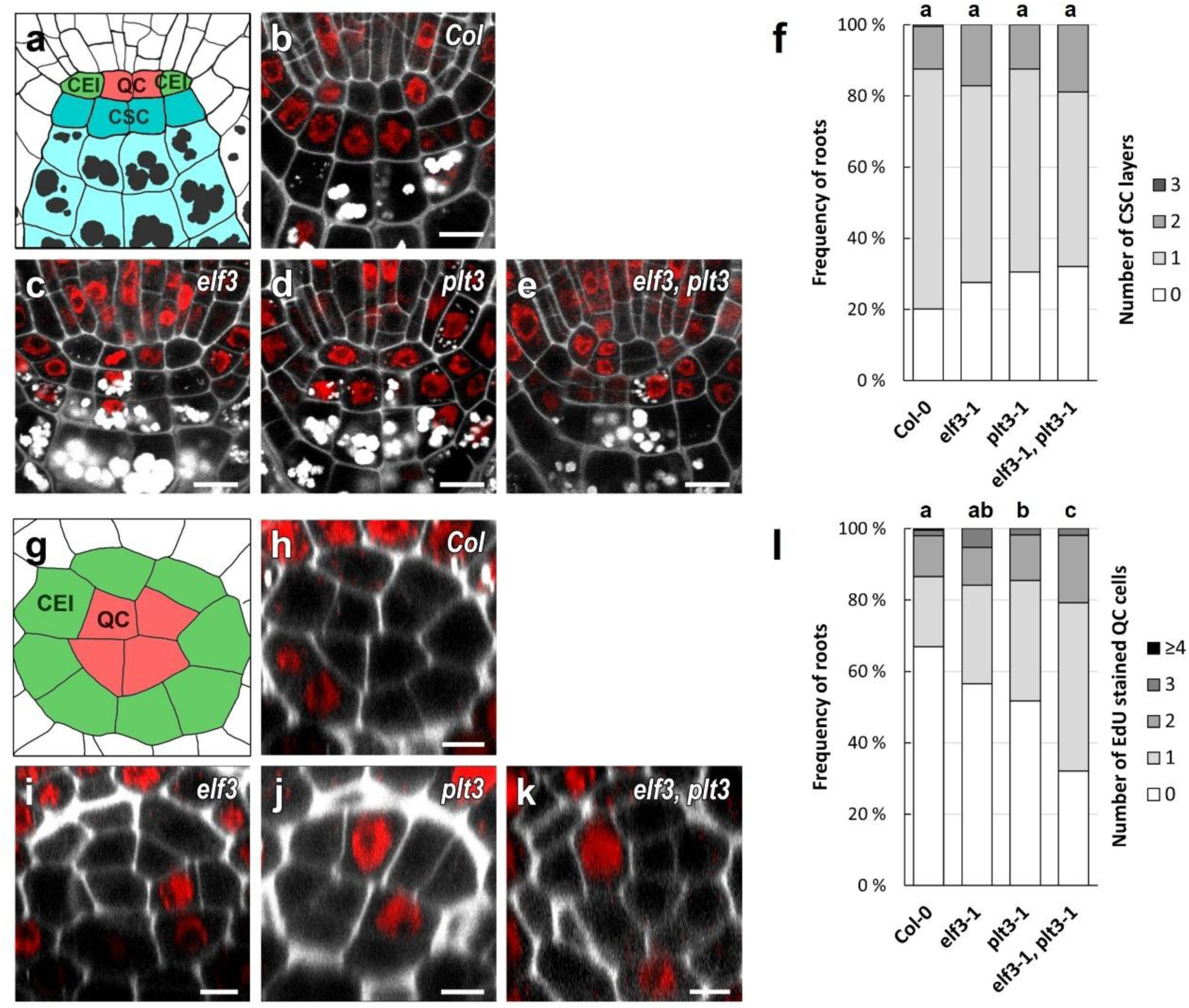
CSC and QC phenotypes of Col-0 wildtype and *elf3-1* and *plt3-1* single and double mutants. **a, g:** Schematic representation of a longitudinal (a) and transversal (g) section of a wild-typic SCN in the *A. thaliana* root. QC= quiescent center (red), CSC= columella stem cells (dark cyan), CEI= cortex-endodermis initials (green). Starch (grey dots) accumulating columella cells are shown in light cyan**. b-e, h-k:** Representative images of the combined mPS-PI (grey) and EdU (red) staining for 24 hours (SCN staining) to analyze the CSC **(b-e)** and QC division **(h-k)** phenotypes within Col-0 and *elf3-1* and *plt3-1* single and double mutants roots. **f:** Analyses of the CSC layers phenotypes, where the frequencies of roots showing 0, 1, 2, or 3 CSC layers are plotted as bar graphs. **l:** Analyses of the QC divisions phenotypes, where the frequencies of roots showing 0, 1, 2, 3 or ≥4 dividing QC cells are plotted as bar graphs. Statistical analysis was performed using a Kruskal-Wallis test followed by Dunn’s multiple comparisons test with Benjamini-Hochberg correction. Groups sharing the same letter are not significantly different (α= 0.05). The number of analyzed roots can be found in Supplementary table 8. SCN= stem cell niche; mPS-PI= modified pseudo-Schiff propidium iodide; EdU= 5-ethynyl-2’-deoxyuridine; scale bars represent 5 µm.

Additionally, QC divisions were assessed in the same roots by counting EdU-stained cells after 24 hours, using z-stacks spanning the root SCN and analyzing the transversal sections. QC cells are known to divide infrequently, with only 20 % undergoing division within 24 hours, roughly half the rate observed in the surrounding initials (Cruz-Ramírez et al. 2013). In this study, 67 % of Col-0 roots showed no QC division, while 32 % had at least one dividing QC cell. However, *elf3-1* and *plt3-1* single mutants displayed significantly higher division frequencies (43 % and 48 %, respectively). Interestingly, the *elf3-1 plt3-1* double mutant showed a further increase to 68 %, suggesting an additive effect and independent roles of ELF3 and PLT3 in the regulation of QC divisions (Figure 2g-l). Statistical analyses confirmed the significant variations among genotypes. These findings indicate an additive effect of ELF3 and PLT3 in controlling QC divisions, suggesting that they act through independent pathways to restrict QC divisions and maintain the SCN homeostasis.

Furthermore, the numbers of QC divisions and CSC layers were negatively correlated (Spearman’s correlation = -0.19, p-value = 1.7 x 10^-5^) in all genotypes. This observation reflects typical SCN balance, where increased differentiation marked by fewer CSC layers trigger compensating QC divisions to maintain SCN homeostasis. This negative correlation was moderate in Col-0, strongest in *elf3-1* and *elf3-1, plt3-1* mutants, and absent in *plt3-1* (Supplementary table 8). Moreover, we performed a permutational multivariate analyses of variance (PERMANOVA) using QC division and CSC layer numbers as joint response variable to assess whether the studied genotypes differed in their overall SCN phenotype. This analysis revealed a significant genotype effect (R^2^=0.02, p-value= 3 x 10^-3^), indicating that ELF3 and PLT3 can significantly alter the SCN overall phenotype.

We plotted 2D histograms showing the number of QC divisions (x-axis) and CSC layers (y-axis), with color-coded root frequencies, following previously described methods (Burkart et al. 2022). Col-0 displayed a defined peak at one CSC layer and no QC divisions (47 %) (Figure 3a). Similar peaks were observed with *elf3-1* and *plt3-1*, but with reduced frequencies (36 % and 31 %, respectively). Also, they shifted towards less CSC layers, i.e., more CSC differentiation, and higher QC divisions, highlighting the correlation between higher root differentiation and reduced quiescence (Figure 3b-c). However, the *elf3-1 plt3-1* double mutant had a wider distribution. It peaked at one CSC layer and one QC division (23 %), and it showed more roots with extreme CSC and QC phenotypes (Figure 3d). These findings suggested that both ELF3 and PLT3 contribute to the SCN homeostasis maintenance. Together, they additively regulate the QC divisions but act interdependently to preserve the CSC layer. Notably, disrupting the QC quiescence influences the CSC differentiation, further supporting that the two proteins promote QC stability and maintain SCN identity (Supplementary table 9).

**Figure 3:**
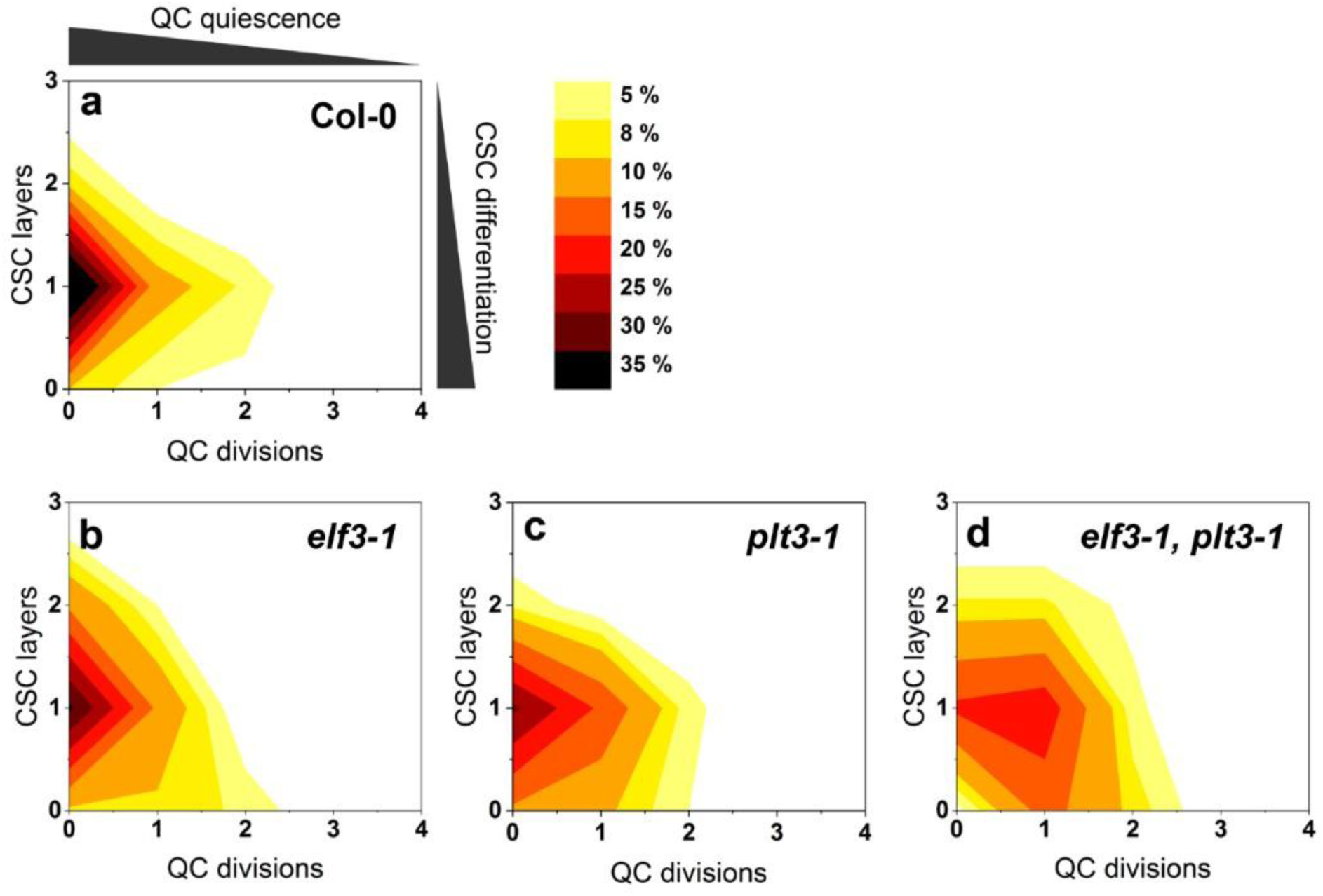
2D visualization of QC divisions and number of CSC layers. **a-d:** The combined results of the SCN staining are shown as 2D plots to visualize the correlation between CSC and QC phenotypes. The number of QC division is plotted on the x-axis, and the number of CSC layers is plotted on the y-axis. The color gradient indicates the frequencies of the observed phenotypes.

### ELF3 and PLT3 proteins localize into subcellular condensates with different behavior

Previous studies have demonstrated that both ELF3 and PLT3 localize to subcellular condensates in *A. thaliana* and in *Nicotiana benthamiana* during transient expression assays (Herrero et al. 2012; Jung et al. 2020; Burkart et al. 2022; Hutin et al. 2023). Based on these findings, we further investigated their subcellular distribution individually and examined their localization and dynamics using different FP tags in a transient *N. benthamiana* expression system under an estradiol-inducible promoter to control expression strength (Bleckmann et al. 2009).

Consistent with previous studies (Jung et al. 2020; Burkart et al. 2022), we observed that both mVenus (mV)-tagged ELF3 and PLT3 fusion proteins localize into subcellular condensates in *A. thaliana* and *N. benthamiana*. In *A. thaliana* roots, both proteins formed condensates, predominantly in the nucleus at room temperature (Figure 1b,c’). In *N. benthamiana*, PLT3 primarily formed nuclear condensates, whereas ELF3 localized within nuclear and cytoplasmic condensates. Since subcellular localization is often associated with differences in condensate material properties and dynamics, we next asked whether nuclear versus cytoplasmic condensates exhibit distinct dynamic behaviors.

To address this, we recorded 3D-time series of cytoplasmic ELF3-mV and nuclear PLT3-mV in *N. benthamiana*, followed by tracking the condensates. ELF3 and PLT3 condensates displayed distinct dynamic behaviors. Here, ELF3 cytoplasmic condensates exhibited fusion and fission events, representative of liquid-like behavior as typical for LLPS (Jung et al. 2020) (Figure 4a-b’’). These condensates were highly mobile, moving over relatively long distances (2.53 ± 4.33 µm) at relatively high speeds (0.74 ± 0.08 µm/s). However, PLT3 condensates were less mobile, as their displacement was limited (0.50 ± 0.16 µm), and they moved approximately ten times slower than ELF3 (0.06 ± 0.001 µm/s) (Figure 4c-f). Additionally, the speed of ELF3 bodies was strongly correlated to their mean intensity (Figure 4g-h), with lower-intensity condensates displaying higher mobility (Supplementary table 14). Higher fluorescence intensities reflect a larger accumulation of fluorescently-labelled proteins. While it is only logical that bigger condensates move slower, yet, in PLT3, this effect was much weaker. Taken together, these observations indicate that PLT3 condensates exhibit lower mobility than ELF3 condensates and suggest that their mobility may be influenced by additional molecular interactions or material properties.

**Figure 4:**
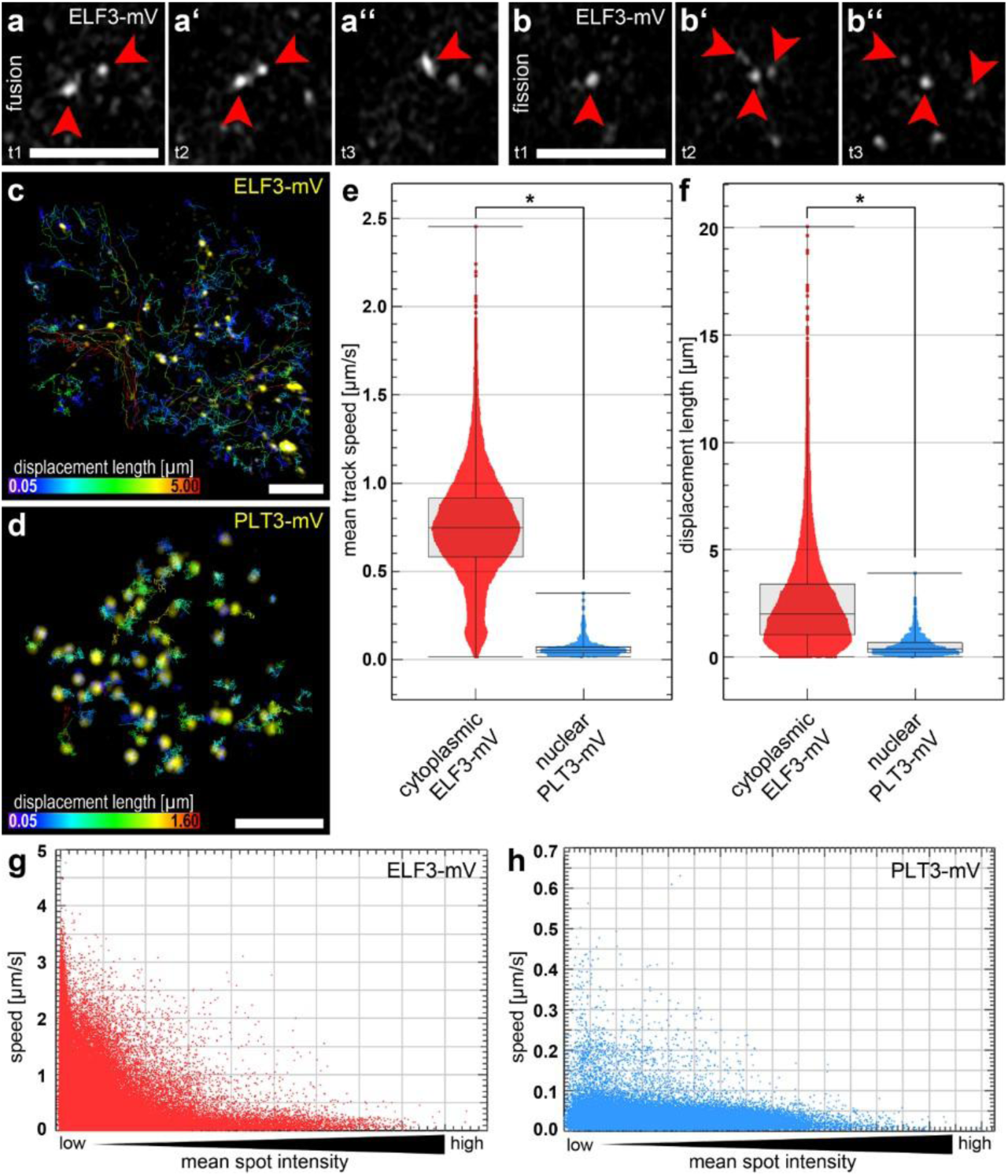
Condensate tracking revealing a higher mobility for ELF3 bodies than PLT3 in *N. benthamiana*. **a-a ’’:** Example for a fusion event of two ELF3-mV condensates. Three time points (t1-t3) are shown. Red arrowheads point at the fusing condensates. **b-b’’:** Example for a fission event of an ELF3-mV condensate. Three time points (t1-t3) are shown. Red arrowheads point at the condensate that undergoes fission into three. **c, d:** Representative MIP-images of tracking experiments of cytoplasmic ELF3-mV **(c, yellow)** and nuclear PLT3-mV **(d, yellow)**. Color-coded lines represent tracks of the mobile condensates, where the color represents the displacement length in µm. **e, f:** Box and scatter plots of the mean track speed **(e)** and the track displacement length **(f)** of cytoplasmic ELF3-condensates (red) and nuclear PLT3-condensates (blue). Asterisks mark statistically significant differences, analyzed with the Kolmogorov-Smirnov-Test and a confidence level of 0.01. The data and number of data points is summarized in Supplementary table 14. Boxes indicate the 25–75% percentile, whiskers show 1.5× the interquartile range, the line marks the median. **g, h:** Scatter plots of the speed of the tracked ELF3-condensates **(g, red)** and PLT3-condensates **(h, blue)** in dependence of their mean mV intensity. Scale bars represent 5 µM. MIP= maximum intensity projection; mV= mVenus.

In PLT3, three intrinsically disordered prion-like domains (PrDs), one located near its N-terminus and two towards its C-terminus are required for its localization in nuclear condensates (Figure 5a) (Burkart et al. 2022). Interestingly, PLT3ΔPrD, a PLT3 variant with mutated PrD domains, still localizes in the nucleus but fails to form condensates (Burkart et al. 2022). As ELF3 also forms subcellular condensates, we analyzed its protein sequence using the web-based PLAAC tool (Lancaster et al. 2014) and confirmed the presence of two PrDs located in the C-terminal part of the ELF3 protein (Figure 5b) (Jung et al. 2020; Hutin et al. 2023). We generated an ELF3ΔPrD variant lacking both C-terminal PrD-enriched regions, where the PrDs were deleted and replaced by a 27 amino acid linker. In the *N. benthamiana* transient expression system, we found that PLT3-mV localized in the nucleus and formed nuclear condensates, while PLT3ΔPrD-mV remained nuclear but showed no condensation, consistent with previous findings (Figure 5c-d) (Burkart et al. 2022). However, ELF3-mV localized in both the nucleus and cytoplasm, forming nuclear and cytoplasmic condensates (Figure 5e). In contrast to PLT3ΔPrD, ELF3ΔPrD-mV still formed condensates at room temperature (Figure 5f). Supporting this, we observed a similar pattern in *A. thaliana* where ELF3ΔPrD-mV expressed in the *elf3-1* mutant background also formed condensates (Supplementary figure 2). Together, these results show that, unlike PLT3, deletion of the PrDs in ELF3 is not sufficient to abolish its condensation in our *in vivo* assays, although these domains have been implicated in ELF3 condensation *in vitro* (Hutin et al. 2023). This suggests that ELF3 condensation can occur through additional protein regions and/or interactions with other cellular compartments.

**Figure 5:**
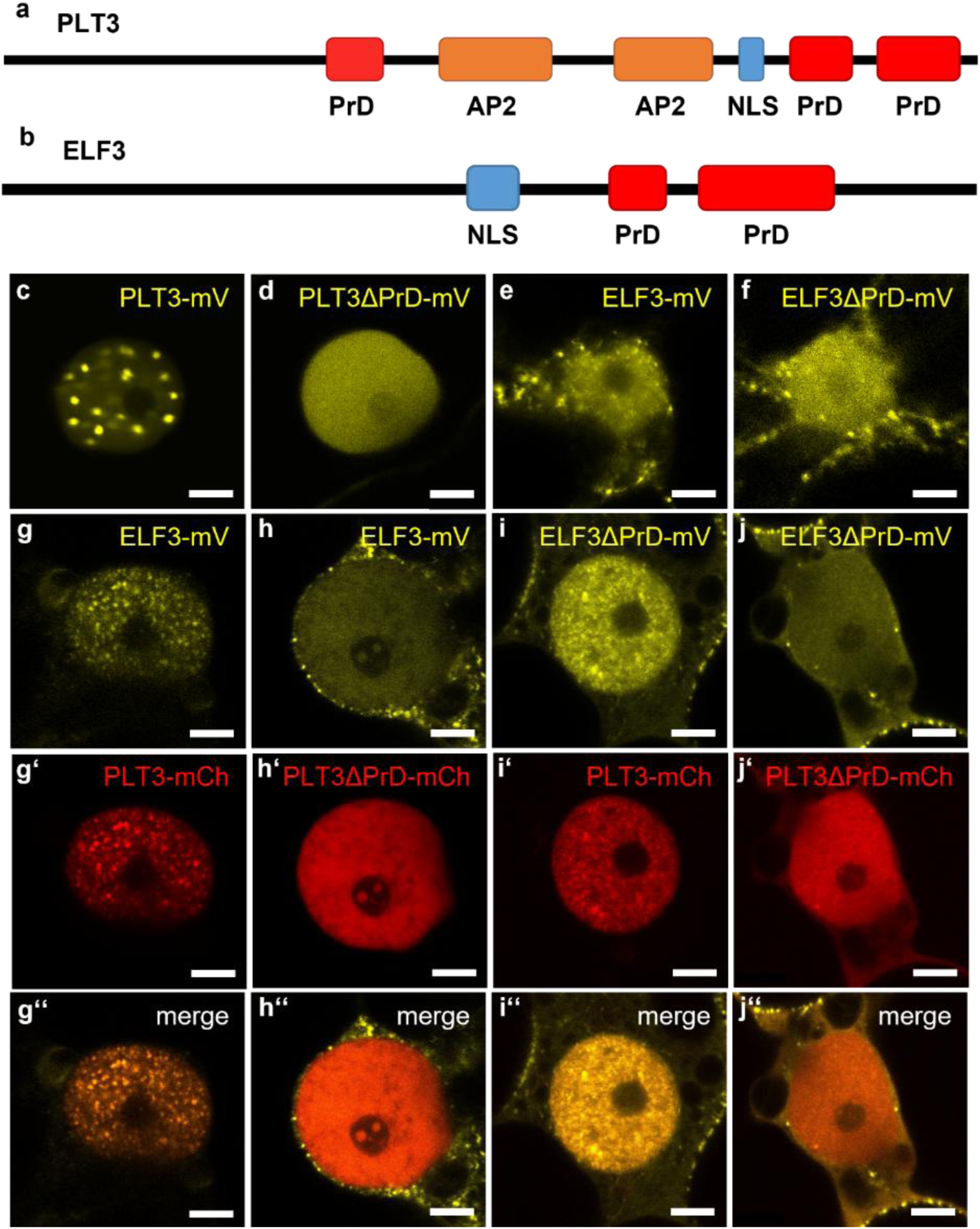
PrD-dependent subcellular (co)-localization of ELF3 and PLT3 in *N. benthamiana*. **a, b:** Schematic representation of the protein domains in PLT3 **(a)** and ELF3 **(b)**. The prion-like domains (PrDs, red) were predicted with a web-based tool (Lancaster et al. 2014). AP2 domains are shown in orange, NLS in blue. **c-f:** Subcellular localization of mV-tagged PLT3 or ELF3-variants. **g-j’’:** Subcellular co-localization of combinations of mV-tagged ELF3-variants (yellow) and mCh-tagged PLT3-variants (red). Scale bars represent 5 µm. mV= mVenus; mCh= mCherry; PrD= Prion-like domain; AP2= APETALA2 domain; NLS= nuclear localization signal.

### PrDs mediate the PLT3-dependent nuclear recruitment of ELF3 and their colocalization into nuclear condensates

Additionally, we co-expressed different variants of mV-tagged ELF3 and mCherry (mCh)-tagged PLT3 to assess their potential colocalization at room temperature. Upon their co-expression, full-length ELF3-mV fully colocalized with PLT3-mCh in nuclear condensates, arguing for a complete translocation of ELF3 from the cytosol to the nucleus in presence of PLT3 (Figure 5g-g’’). Interestingly, when ELF3-mV was co-expressed with PLT3ΔPrD-mCh, the nuclear recruitment of ELF3 was abolished and no nuclear condensates were formed (Figure 5h-h’’). In contrast, the co-expression of ELF3ΔPrD-mV with the full length PLT3-mCh resulted in nuclear localization and condensate formation comparable to that of full-length ELF3 (Figure 5i-i’’). When PrDs from both proteins were mutated, nuclear condensates were no longer observed, while ELF3 still displayed cytoplasmic condensates (Figure 5j-j’’). These results suggest that ELF3 can be recruited to the nucleus in a PLT3-dependent manner, where the PrDs of PLT3, but not those of ELF3, are important for ELF3 nuclear import. This recruitment seems to be associated with increased ELF3 nuclear accumulation and reduced cytoplasmic signal.

To quantify the PLT3-dependent nuclear import of ELF3, we measured the mean fluorescence intensities of mV-tagged ELF3 variants, both alone or in combination with mCh-tagged PLT3 variants, in the nucleus and cytoplasm. In this experiment, a change in the mVenus intensity ratio (nucleus/cytoplasm) reflects the nuclear translocation of ELF3. Initially, no significant change in the nuclear/cytoplasmic distribution of ELF3 was observed between ELF3-mV (ratio= 1.67 ± 0.72) and ELF3ΔPrD-mV (ratio= 2.47 ± 1.47). However, the intensity ratio upon ELF3-mV co-expression with PLT3-mCh increased significantly about six-fold (ratio= 11.86 ± 8.54), verifying the PLT3-dependent nuclear import of ELF3. Interestingly, this import was impaired upon PrD deletion in PLT3, as the ratio decreases in ELF3-mV and PLT3ΔPrD-mCh co-expression (ratio= 2.14 ± 1.17) (Figure 6, Supplementary table 10).

**Figure 6:**
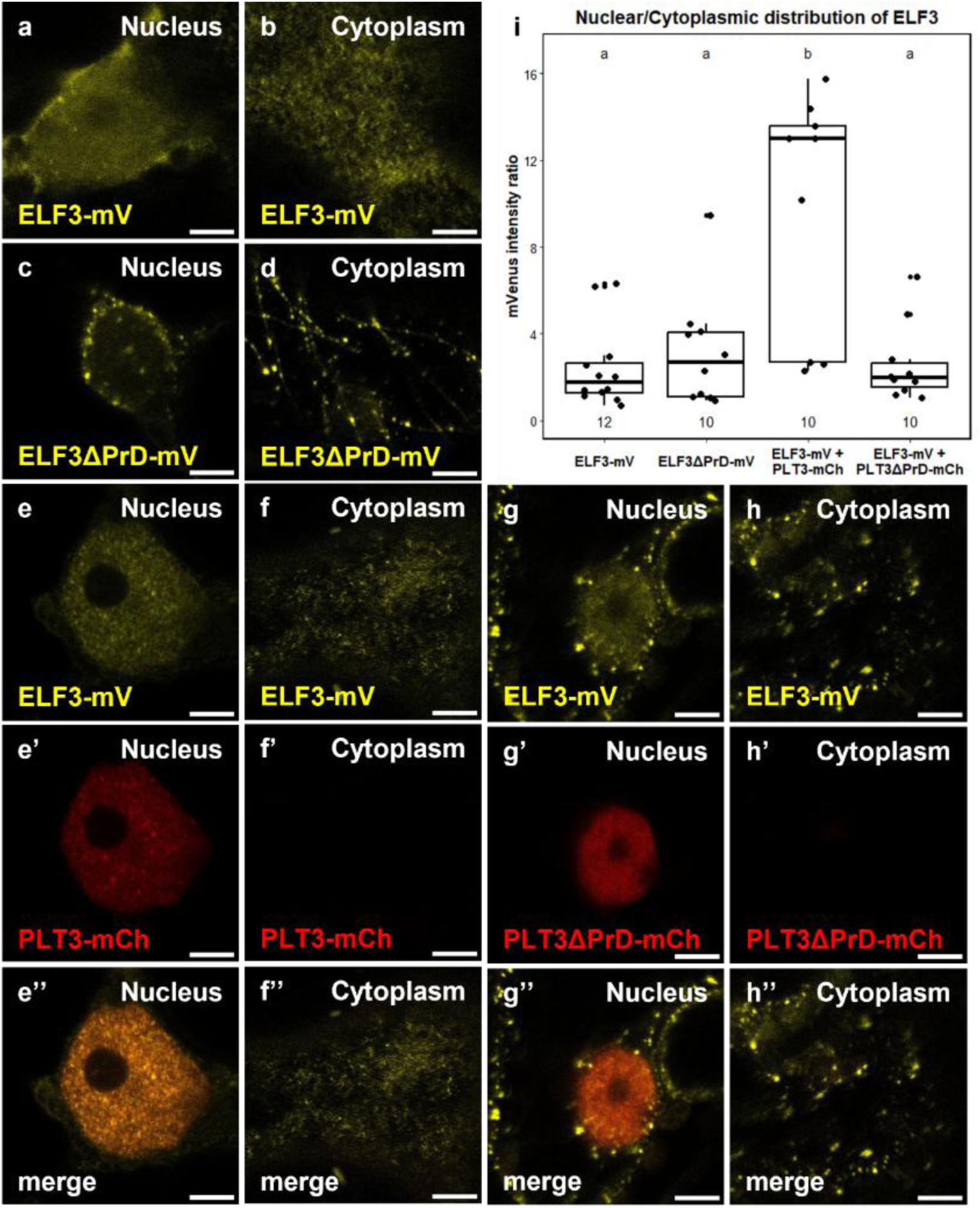
PLT3-dependent nuclear recruitment of ELF3 by PLT3 PrDs. **a-d:** Representative images of nuclear (left) and corresponding cytoplasmic (right) localization of mV-tagged ELF3-variants (yellow) in *N. benthamiana*. **e-h’’:** Representative images of nuclear (left) and corresponding cytoplasmic (right) localization of ELF3-mV (yellow) upon co-expression with mCh-tagged PLT3-variants (red) in *N. benthamiana*. **i:** Scatter and box plots presenting mVenus intensity ratios. Each dot represents a single measurement, boxes indicate the 25–75% percentile, whiskers show 1.5 x the interquartile range, the line marks the median, and the square denotes the mean. Replicate numbers are shown below each group. Statistical analysis was performed using a Kruskal-Wallis test followed by Dunn’s multiple comparisons test with Benjamini-Hochberg correction. Groups sharing the same letter are not significantly different (α= 0.05). Scale bars represent 5 µm. mV= mVenus; mCh= mCherry.

Therefore, *in vivo*, ELF3 is imported to the nucleus and recruited to nuclear condensates in a PLT3-dependent manner, mediated by the PLT3 PrDs. Given the similarities in amino acid composition between the PrD of ELF3 and the second PrD of PLT3, namely the presence of polyQ stretches (Supplementary figure 3a), we investigated whether the second PrD of PLT3 can multimerize or drive phase separation *in vitro* in a similar manner as previously observed in vitro for ELF3 PrD (Hutin et al. 2023). Structural studies using small angle X-ray scattering (SAXS) analysis showed that, while the ELF3 PrD forms a large globular multimer (Hutin et al. 2023) (Supplementary figure 3b), the PLT3 PrD2 is monomeric, as confirmed by Size-Exclusion Chromatography-Multi-Angle Laser Light Scattering (SEC-MALLS) (Supplementary figure 3c). Furthermore, PLT3 PrD2, based on Kratky analysis, exhibited a plateau at high q, indicative of an intrinsically disordered protein (Supplementary figure 3b), in contrast to ELF3 PrD, which exhibits a bell-shaped profile characteristic of a globular species (Hutin 2023). We investigated the ability of PLT3 PrD2 to form condensates *in vitro*, however, we were not able to trigger the condensation of PLT3 PrD2 alone under our tested conditions, while ELF3 PrD easily underwent phase separation. Attempts to form a complex between ELF3 PrD and PLT3 PrD2 were not successful and we could not detect direct interaction between these protein domains, suggesting that the interaction requires additional regions of the proteins or cofactors *in vivo*.

### The *in vivo* ELF3-PLT3 interaction is PrD-mediated

The ELF3-PLT3 co-localization in nuclear condensates observed *in vivo* prompted us to investigate their direct physical interaction. Thus, we performed Förster Resonance Energy Transfer -Fluorescence Lifetime Imaging Microscopy (FRET-FLIM) to test their *in vivo* protein-protein interaction (PPI) and whether it is mediated by their PrDs (Strotmann and Stahl 2022). In plants, FRET-FLIM is now commonly used to study PPIs with high spatiotemporal resolution resulting in a decrease of the donoŕs fluorescence lifetime in case of PPI (Eljebbawi et al. 2025; Strotmann et al. 2025).

We used mV-tagged ELF3 variants as donors (ELF3-mV and ELF3ΔPrD-mV) and mCh-tagged PLT3 variants as acceptors (PLT3-mCh and PLT3ΔPrD-mCh) (Figure 7, Supplementary table 15). As negative controls, free-mCh was co-expressed with the respective donors. Initially, the donor-only fluorescent lifetime was measured for ELF3-mV (τ= 2.99 ± 0.06 ns) and ELF3ΔPrD-mV (τ= 3.03 ± 0.04 ns). The negative controls did not show any significant decrease in mVenus fluorescence lifetime compared to the corresponding donor-only samples (τ= 2.99 ± 0.04 ns for ELF3-mV + free-mCh and τ= 3.00 ± 0.02 ns for ELF3ΔPrD-mV + free-mCh).

**Figure 7:**
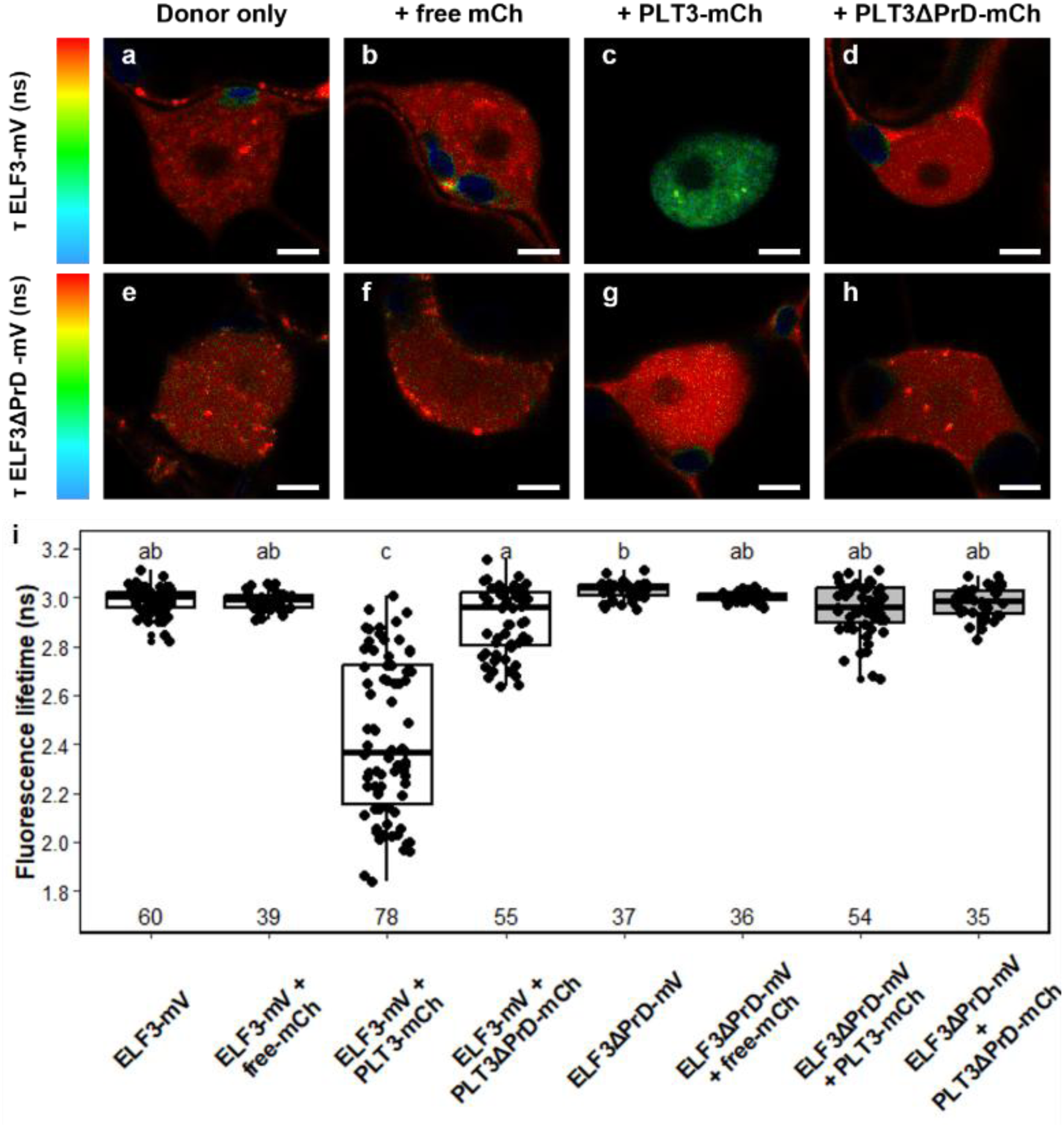
The PrD-mediate interaction of ELF3 and PLT3. **a-h:** Representative images of mVenus fluorescence lifetime (ns) of mV-tagged ELF3-variants alone or co-expressed with free mCherry (mCh) and mCh-tagged PLT3-variants. The mV fluorescence lifetime is color-coded from blue (shorter lifetime) to red (longer lifetime). Dark blue structures are chloroplasts and were not included in data analysis. Scale bars represent 5 µm. **i:** Scatter and box plots presenting fluorescence lifetime (ns). Each dot represents a single measurement, boxes indicate the 25–75% percentile, whiskers show 1.5 x the interquartile range, the line marks the median, and the square denotes the mean. Replicate numbers are shown below each group. Replicate numbers are shown below each group. Statistical analysis was performed using a Kruskal-Wallis test followed by Dunn’s multiple comparisons test with Benjamini-Hochberg correction. Groups sharing the same letter are not significantly different (α= 0.05).

Upon the co-expression of ELF3-mV and PLT3-mCh, we noticed a significant reduction in the mVenus fluorescence lifetime (τ= 2.69 ± 0.15 ns), indicating a direct interaction between the full-length ELF3 and PLT3 proteins. Additionally, pull-down experiments of recombinant full length ELF3 and PLT3 proteins confirmed their direct interaction (Supplementary figure 4). However, *in vivo,* no significant fluorescent lifetime reductions were detected when the PrDs of either or both proteins were mutated. Here, the fluorescent lifetimes were 2.91 ± 0.14 ns for ELF3-mV co-expressed with PLT3ΔPrD-mCh, 2.95 ± 0.1 ns for ELF3ΔPrD-mV co-expressed with PLT3-mCh, and 2.98 ± 0.07 ns for the double-mutated pair ELF3ΔPrD-mV and PLT3ΔPrD-mCh (Figure 7, Supplementary table 15). These results demonstrated that the deletion of the PrD domains of either ELF3 or PLT3 was sufficient to abolish the detectable FRET signal, indicating that the direct ELF3-PLT3 interaction detected *in vivo* requires these regions. Notably, ELF3ΔPrD remained nuclear and could still localize to nuclear condensates, indicating that co-localization within condensates does not necessarily correspond to detectable direct ELF3-PLT3 interaction. This further suggests that PLT3-mediated nuclear import of ELF3 may occur indirectly and/or require additional interaction partners.

### PIFs facilitate ELF3 nuclear translocation and its association with PLT3

In our transient expression experiments, we used *N. benthamiana* leaves as a tool to examine the studied *A. thaliana* proteins in a plant cellular environment. This method has been widely reported to provide reliable *in vivo* subcellular localization and PPI measurements (Sparkes et al. 2006; Cody et al. 2023; Beritza et al. 2024). However, the presence of other phylogenetically-relevant plant proteins could bias our observations. Therefore, we used mammalian Human epithelial cells HEp-2 (epithelial larynx carcinoma, ATCC-CCL-23) as a non-plant orthogonal system for transient protein expression. Here, we asked whether we would observe similar localization of PLT3 and ELF3 in this environment in the absence of other plant-specific factors. Similar to *N. benthamiana*, PLT3-mV localized in the nucleus and formed nuclear condensates (Figure 8a). As for ELF3-mV, it predominantly localized in the cytoplasm, where it formed cytoplasmic condensates, but it was observed to a much lesser extent in HEp-2 nuclei compared to *N. benthamiana* (Figure 8b). Interestingly, when we co-expressed ELF3-mV with mRuby2 (mRb)-tagged PLT3 in HEp-2 cells, ELF3 was not recruited to the nucleus but remained predominantly in the cytoplasm, and the two proteins did not colocalize, in contrast to our observation in *N. benthamiana* (Figure 8e-e’’). This finding suggests that PLT3 alone is insufficient to efficiently recruit ELF3 to nuclear condensates in Hep-2 cells.

**Figure 8:**
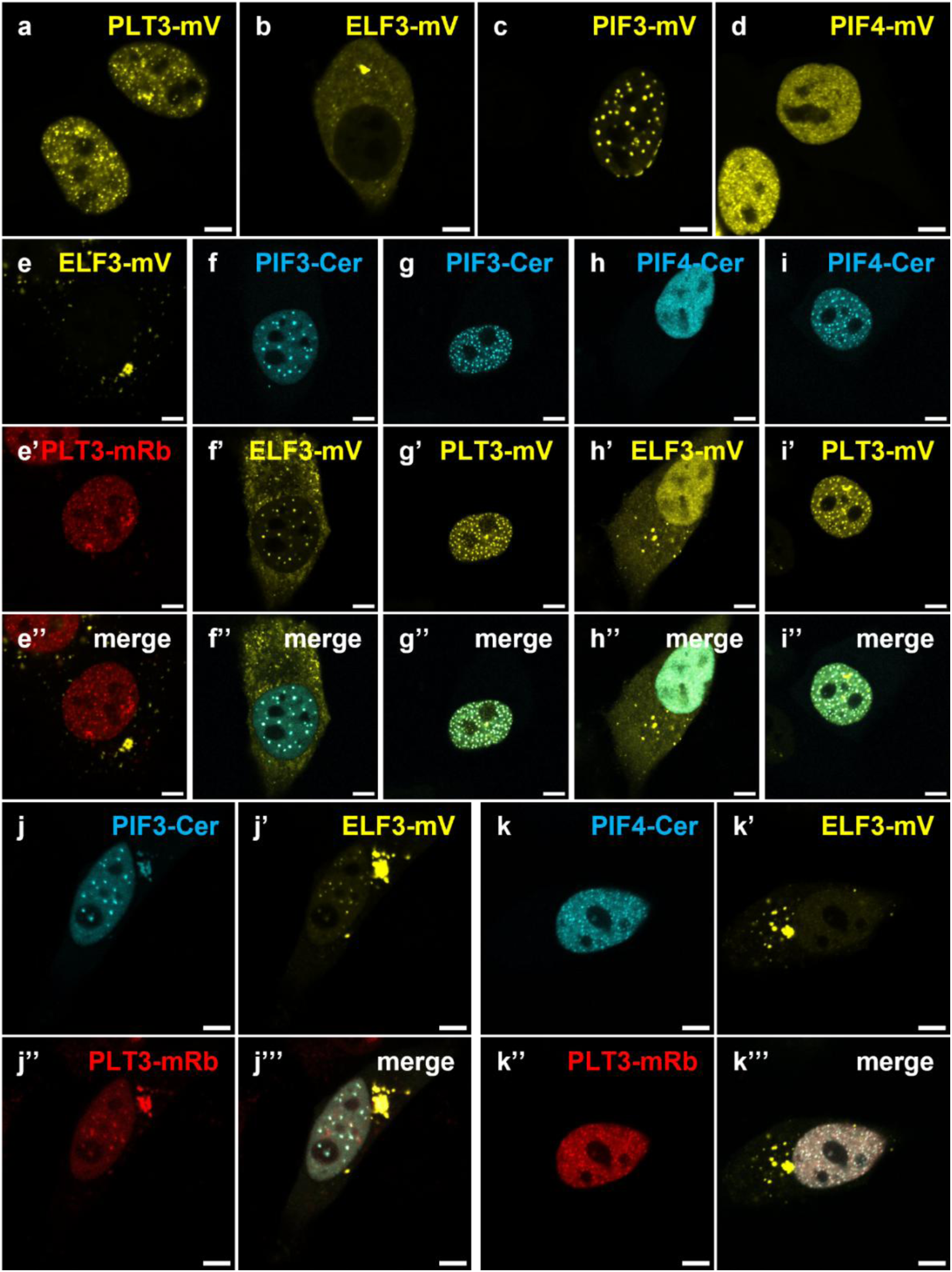
PIF3 and PIF4 import ELF3 to the nucleus and colocalize with PLT3 in HEp-2 cells. **a-d:** Subcellular localization of mV-tagged PLT3, ELF3, PIF3, and PIF4 in HEp-2 cells. **e-e’’:** ELF3-mV (yellow) localizes to cytoplasmic condensates when co-expressed with PLT3-mRb (red), which forms nuclear condensates. **f-i’’:** ELF3-mV (yellow) and PLT3-mV colocalize with PIF3-Cer (cyan) and PIF4-Cer (cyan) in the nucleus within condensates. **j-k’’’:** FP-tagged ELF3 (yellow) and PLT3 (red) colocalize in the nucleus with PIF3-Cer and PIF4-Cer. Scale bars represent 5 µm. mV= mVenus; mRb= mRuby2; Cer= Cerulean.

Previous experiments in mammalian expression systems, i.e., HEp-2 cells, showed that PIF3 can mediate the nuclear import of interacting proteins. For instance, during light response, PIF3 binds and shuttles the photoactivated PHYTOCHROME B (phyB) to the nucleus, where they colocalize in nuclear condensates (Beyer et al. 2015). This led us to inquire whether PIFs might facilitate ELF3 nuclear shuttling and promote the co-localization of ELF3 and PLT3 in the nucleus. Since ELF3 is known to interact with various PIF proteins, such as PIF4 (Nieto et al. 2015), we hypothesized that PIFs potentially facilitate ELF3 nuclear recruitment and thereby promote its association with PLT3. First, we confirmed direct interactions of ELF3, PLT3, PIF3 and PIF4 by immunoprecipitation of recombinant proteins (Supplementary figure 4). These experiments established that ELF3 and PLT3 are capable of directly interacting in the absence of PIFs.

Then we used the HEp-2 system to study the localization of the different components. Both PIF3-mV and PIF4-mV localized in the nucleus and formed nuclear condensates, where PIF3 had generally larger condensates than PIF4 (Figure 8c-d). Notably, when ELF3 was co-expressed with Cerulean (Cer)-tagged PIF3, it maintained its cytoplasmic localization but, remarkably, also colocalized with PIF3 in nuclear condensates (Figure 8f-f’’). This observation suggests that PIF3 can facilitate ELF3 nuclear recruitment. Additionally, PIF3-Cer colocalized with PLT3-mV in nuclear condensates (Figure 8g-g’’). We obtained comparable results when we co-expressed PIF4-Cer with ELF3-mV or PLT3-mV (Figure 8h-i’’), indicating that PIF4 can also facilitate ELF3 nuclear recruitment and co-localize with PLT3 in nuclear condensates. Furthermore, the co-expression of ELF3-mV and PLT3-mRb with PIF3-Cer (Figure 8j-j’’’) or PIF4-Cer (Figure 8k-k’’’) resulted in the colocalization of all three proteins to nuclear condensates. Altogether, our results show that PIF3 and PIF4 facilitate the nuclear recruitment of ELF3 and thereby promote its co-localization with PLT3 in nuclear condensates. Importantly, our FRET-FLIM and recombinant protein interaction assays demonstrate that ELF3 and PLT3 can directly interact independently of PIF3 or PIF4. Thus, PIF3 and PIF4 are not required for the direct ELF3-PLT3 interaction but may facilitate their encounter in the nucleus by promoting ELF3 nuclear recruitment and co-localization with PLT3.

### PIF3 and PIF4 colocalize and interact with ELF3 and PLT3 *in planta*

After observing that ELF3, PLT3, PIF3, and PIF4 colocalize in HEp-2 cells, we aimed to verify whether this localization is conserved in plants. Therefore, we performed transient expression experiments in *N. benthamiana* using FP-tagged proteins to examine their *in planta* localization and interactions. Both mV-tagged PIF3 and PIF4 localized in the nucleus where they formed bright condensates, similar to our observation in HEp-2 cells (Figure 9a-b). Upon the co-expression of either PIF3-mCh or PIF4-mCh with PLT3-mV, both proteins colocalized in nuclear condensates (Figure9c-d’’). Additionally, when either PIF3-mCh or PIF4-mCh were co-expressed with PLT3-Cer and ELF3-mV, the three proteins colocalized within the same nuclear condensates (Figure9e-f’’’).

**Figure 9:**
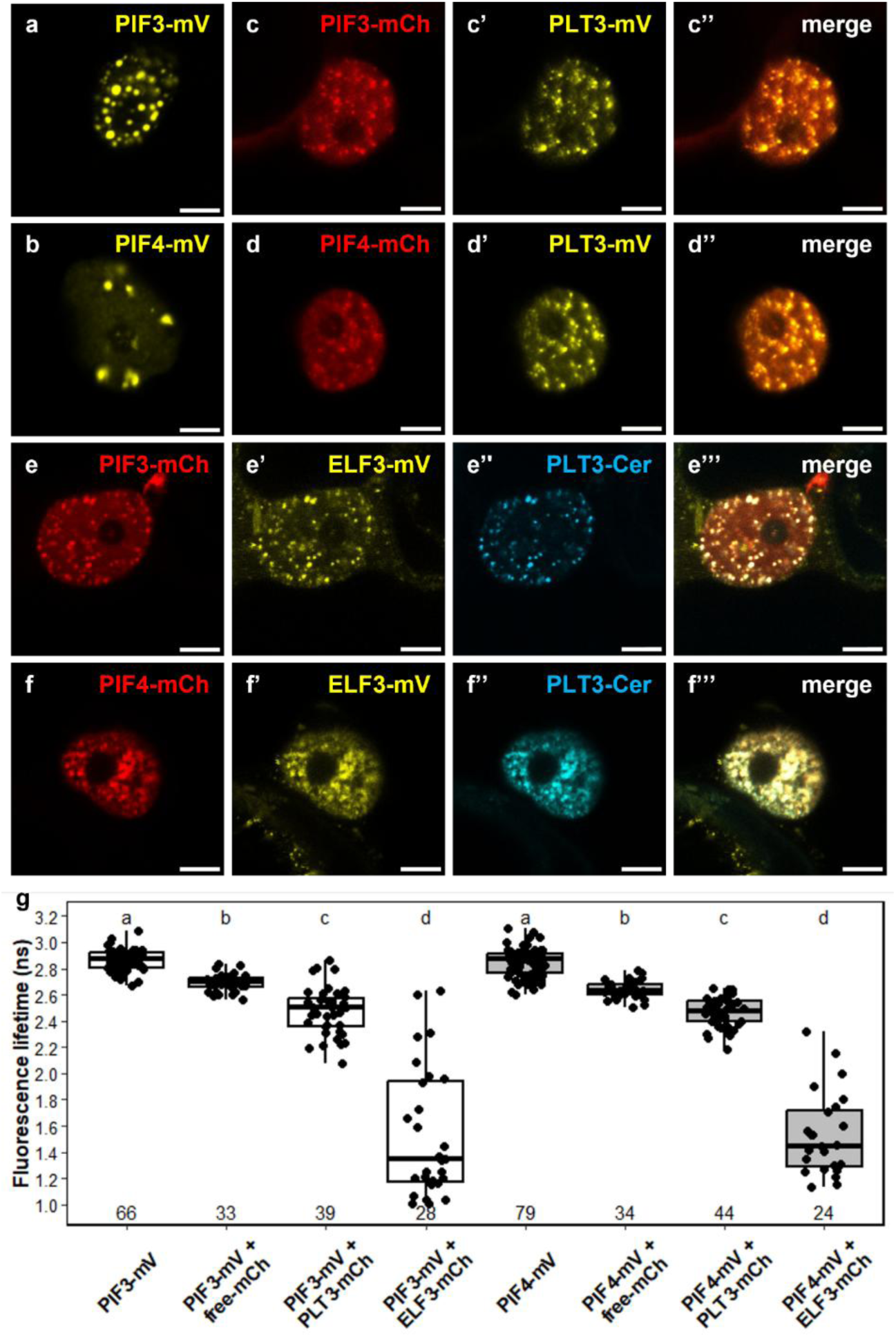
ELF3 and PLT3 colocalize and interact with PIF3 and PIF4 in plants. **a-b :** Subcellular localization of mVenus-tagged PIF3 and PIF4 in nuclear condensates. **c-d’’:** Co-localization of mCherry-tagged PIF3 and PIF4 (red) to nuclear condensates with PLT3-mVenus (yellow). **e-f’’’:** Co-localization of PIF3-mCh or PIF4-mCh (red) with ELF3-mV (yellow) and PLT3-Cer (blue) to nuclear condensates. **g:** Scatter and box plots presenting fluorescence lifetime (ns). Each dot represents a single measurement, boxes indicate the 25–75% percentile, whiskers show 1.5 x the interquartile range, the line marks the median, and the square denotes the mean. Replicate numbers are shown below each group. Statistical analysis was performed using a Kruskal-Wallis test followed by Dunn’s multiple comparisons test with Benjamini-Hochberg correction. Groups sharing the same letter are not significantly different (α= 0.05). mV= mVenus; mCh= mCherry; Cer= Cerulean.

As all four proteins colocalize, we aimed to examine their PPI potential. Accordingly, we conducted FRET-FLIM measurements to verify and further examine the potential interactions of PIF3 and PIF4 together with PLT3 and ELF3. PIF3-mV and PIF4-mV were used as donor-only sample, and their measured fluorescence lifetimes were τ= 2.87 ± 0.08 ns and τ= 2.85 ± 0.11 ns, respectively (Supplementary table 16). As negative controls, PIF3 and PIF4 were respectively co-expressed with free-mCh, where their lifetimes recorded τ= 2.69 ± 0.07 and τ= 2.64 ± 0.06. When co-expressed with PLT3-mCh, the mVenus fluorescence lifetimes significantly decreased compared to both the donor-only samples and negative controls, to τ= 2.48 ± 0.18 ns for PIF3-mV and τ= 2.47 ± 0.11 ns for PIF4-mV. Furthermore, the mVenus fluorescence lifetime was τ= 1.54 ± 0.5 ns for PIF3-mV and τ= 1.53 ± 0.31 ns for PIF4-mV, when co-expressed with ELF3-mCh (Figure 9g, Supplementary table 16). Together, these findings provide the first evidence that PLT3 interacts with both PIF3 and PIF4 *in vitro* (supplementary Figure 4) and *in planta* (Figure 9), suggesting a potential role for the PIF proteins in root SCN maintenance. Additionally, they validate *in vivo* the interaction between ELF3 and PIF4, and confirm, for the first time, the direct interaction between ELF3 and PIF3.

## Discussion

In this study, we uncovered a new role of ELF3 in SCN homeostasis maintenance. An ELF3 reporter line showed that ELF3 is actively expressed in the RAM, including the SCN. Although ELF3 expression in the root has been reported previously (Hicks 2001; Yazdanbakhsh et al. 2011; Jung et al. 2020; Nomoto et al. 2026), its expression within the SCN has not yet been described. Together with PLT3, ELF3 plays a critical role in maintaining the SCN by regulating QC di visions. The QC cells are a long-term reservoir of stem cells with low division rates, surrounded by proliferating initials (Cruz-Ramírez et al. 2013; Vilarrasa-Blasi et al. 2014). PLTs are known master regulators of QC quiescence and CSC maintenance, interconnecting QC divisions and CSC differentiation (Aida et al. 2004; Galinha et al. 2007; Mähönen et al. 2014; Burkart et al. 2022). *PLT1*, *PLT2*, *PLT3*, and *PLT4* are expressed in the *A. thaliana* RAM. They are induced by auxin and form protein gradients that peak in the QC. Functionally, PLTs regulate SCN patterning in a morphogen-like manner, i.e., their gradient distribution determines the cell fate. High PLT levels maintain stem cell identity, while lower levels promote differentiation (Aida et al. 2004; Galinha et al. 2007). Consistent with our findings, PLT3 is known to be concentrated in the QC and CSC layer, where it colocalizes and interacts with WOX5 to maintain QC and CSCs identities (Burkart et al. 2022). Here, we describe for the first time that ELF3 is expressed in the *Arabidopsis* root SCN and additively controls QC divisions, suggesting that it acts in a pathway partially independent of PLT3. In addition to its previously-established roles as a thermosensor and circadian regulator (Covington 2001; Liu 2001; Nusinow et al. 2011; Jung et al. 2020), our results indicate that ELF3 restricts excessive QC divisions, which is pivotal for ensuring a balance between stem cell division and differentiation, therefore maintaining the SCN homeostasis.

ELF3, a transcriptional repressor, has been previously shown to localize into condensates at elevated temperatures, preventing its binding to its target genes, thus lifting the transcriptional repression of flowering-related genes (Jung et al. 2020). Recently, several plant studies highlighted such a role of nuclear condensates in transcriptional regulation, such as the SUPPRESSOR OF MORE AXILLARY GROWTH 2-LIKE 7 (SMXL7), a strigolactone signaling repressor, whose condensates repress transcription by sequestering TFs from promoters (Li et al. 2025), while the transcriptional regulator SEUSS (SEU) undergoes condensation in response to hyperosmotic stress forming hubs to recruit TFs leading to the activation of stress-responsive genes (Wang et al. 2022). Thus, condensates can perform different functional roles, acting to either sequester and inactivate TFs or increase their local concentration and augmenting their transcriptional activity.

Our transient expression experiments performed in *N. benthamiana* demonstrated that ELF3 forms nuclear and cytoplasmic condensates, and it colocalizes with PLT3 in the nucleus. The differences in their condensate mobility suggests that they have distinct biophysical properties, with ELF3 being more dynamic while PLT3 being less mobile. Although the molecular basis for these differences remains unclear, they may reflect differences in condensate composition or interactions with other cellular components. Moreover, these findings are consistent with previous evidence on the liquid-like nature of ELF3-condensates. ELF3 has been shown to form *in vitro* and *in vivo* liquid droplets in response to temperature and pH (Hutin et al. 2023). Also, it recovers during *in vivo* FRAP experiments, a characteristic of liquid-like behavior (Jung et al. 2020; Hutin et al. 2023). Interestingly, PLT3 enhances the nuclear localization of ELF3 when co-expressed in *N. benthamiana*, where this nuclear recruitment requires the PrDs of PLT3. In contrast, PrD-deletion of ELF3 did not abolish its nuclear localization or condensate formation. Furthermore, ELF3 and PLT3 can directly interact, and our FRET-FLIM measurements showed that the PrD deletion of either protein abolished the detectable FRET signal. Thus, the PrDs of both ELF3 and PLT3 contribute to their direct *in vivo* interaction, whereas ELF3 nuclear localization and condensate formation can occur independently of its own PrD domains.

The distinction between nuclear recruitment and direct interaction is further supported by our experiments in HEp-2 cells. In this system, PLT3 alone was insufficient to efficiently recruit ELF3 to the nucleus, despite the ability of ELF3 and PLT3 to directly interact. In contrast, PIF3 and PIF4 facilitated ELF3 nuclear recruitment and resulted in the co-localization of ELF3 and PLT3 within nuclear condensates. Thus, ELF3 and PLT3 possess an intrinsic capacity for direct interaction, but their encounter within the nucleus may be facilitated by additional factors such as PIF3 and PIF4. PIF3 and PIF4 therefore emerge as potential regulators of ELF3 nuclear recruitment and its spatial association with PLT3. PIFs are key developmental regulators, as PIF4 and PIF5 promote hypocotyl growth, and their transcription is repressed by the EC in the evening (Nusinow et al. 2011). PIF3 also promotes hypocotyl growth in a circadian-dependent manner, and it contributes to the inhibition of root growth in response to nitric oxide (Soy et al. 2012; Bai et al. 2014). Importantly, several PIFs are expressed in roots including PIF1, PIF3, PIF4, and PIF5, where they regulate root growth in response to hormonal signaling and environmental cues such as gravity (Ruiz-Sola et al. 2014; Yang et al. 2020). For instance, they are involved in root penetration into the soil where PIF3 stabilizes the receptor kinase FERONIA (FER) in the root cap cells and regulates the expression of the mechanosensitive ion channel *PIEZO*, required for mechanical stress response (Xu et al. 2024). Furthermore, PIF3 inhibits primary root growth in response to nitric oxide (NO) (Bai et al. 2014).

Our findings further identify physical interactions between PLT3 and PIF3/PIF4 and between ELF3 and PIF3/PIF4. In HEp-2 cells, PIF3 and PIF4 were sufficient to facilitate ELF3 nuclear recruitment, and in *N. benthamiana*, all three proteins could be detected within the same nuclear condensates. These observations provide a potential molecular link between PIF-dependent environmental signaling and PLT3-containing nuclear condensates in the root SCN. In addition, we validate the previously reported interaction between ELF3 and PIF4 and provide evidence for a direct interaction between ELF3 and PIF3. It has recently been reported that PIF3 and PIF4 interact, as demonstrated by co-immunoprecipitation in *A. thaliana*, *in vitro* yeast two-hybrid (Y2H) screening, and *in vivo* bimolecular fluorescence complementation (BiFC) assay. Their interaction is required to regulate growth during thermomorphogenesis in *A. thaliana* (Das et al. 2025). Additionally, it has been demonstrated that the EC directly binds to the promoter of PIF4 and PIF5, repressing these targets, thus functionally linking ELF3 to PIF4 and PIF5 (Nusinow et al. 2011). Moreover, a previous study has shown a direct physical interaction between ELF3 and PIF4 through the PIF4 bHLH domain and ELF3 C-terminal regions using Y2H and BiFC (Nieto et al. 2015).

Previously, we proposed that PLT3 and WOX5 regulate the root SCN by forming nuclear condensates, providing a fast and reversible regulatory mechanism for cell fate determination in the RAM (Burkart et al. 2022). In this study, we extend this model by including ELF3 and PIFs as additional regulatory components that may link environmental signaling to SCN homeostasis (Figure 10). ELF3 and PIFs are known integrators of a circadian, light, and temperature-dependent signaling, as well as growth-related processes such as photosynthesis and hormonal signaling (Nusinow et al. 2011; Huang et al. 2016; Chow et al. 2012; Ezer et al. 2017; Silva et al. 2020; Jung et al. 2020). In our model, ELF3 may act as a dynamic sensor of environmental signals in the root SCN. In response to specific environmental stimuli, ELF3 condensates dynamically redistribute between the cytoplasm and the nucleus, facilitated by PIFs that are expressed throughout the RAM (Figure 10) (Birnbaum et al. 2003; Nawy et al. 2005). Within the nucleus, ELF3 can directly interact with PLT3, mediated by the PrDs of both proteins. PIF3 and PIF4 may further facilitate the co-localization of ELF3 and PLT3 within nuclear condensates through their interactions with both proteins. Together, these interactions may provide a mechanism to integrate environmental signaling with PLT3-dependent regulation of SCN homeostasis, potentially allowing root stem cell activity to respond dynamically to changing environmental conditions. While our data support this model, the precise underlying molecular mechanisms governing condensate assembly, dynamics, and downstream regulatory output remain to be elucidated.

**Figure 10:**
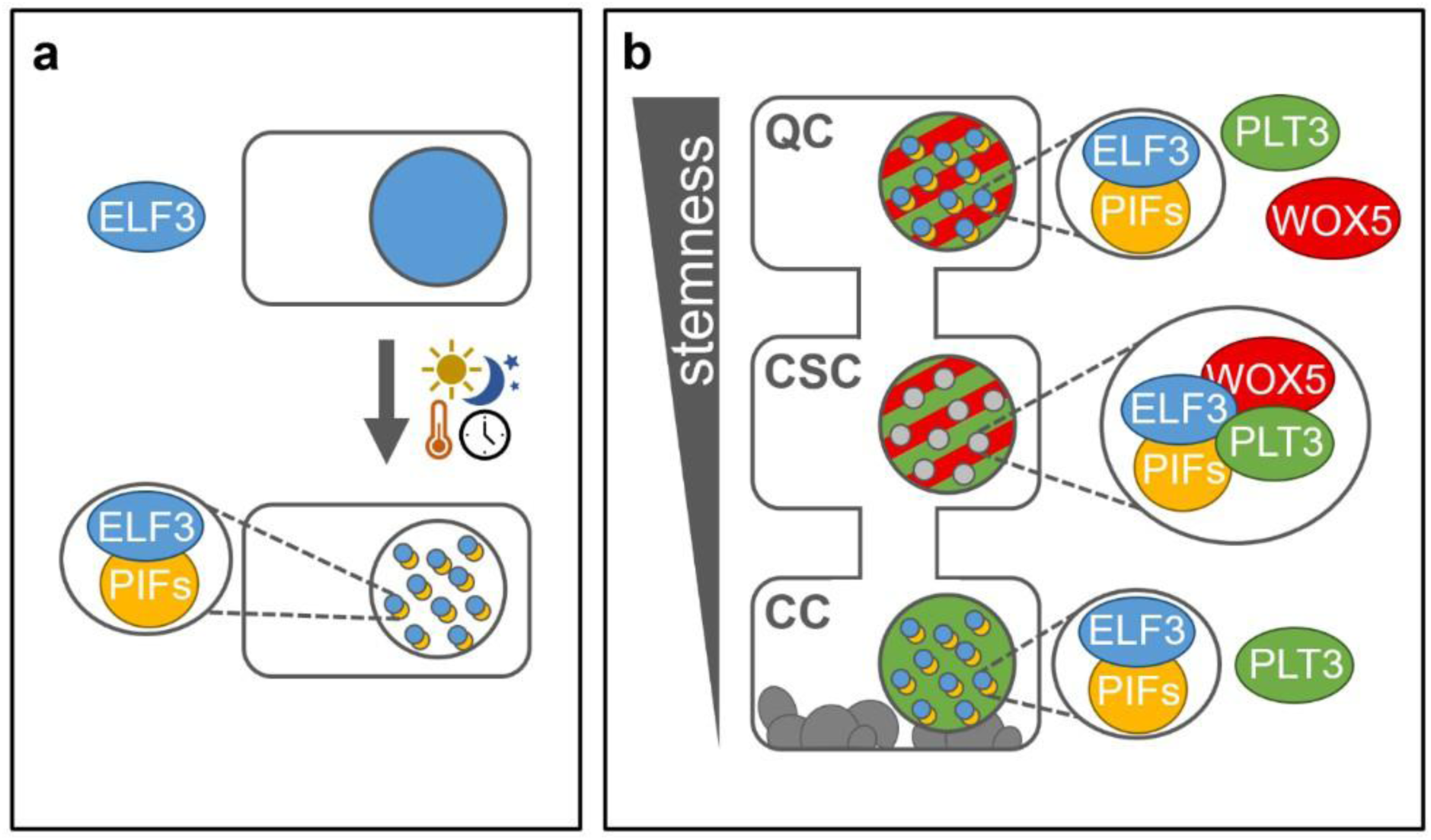
Model of *A. thaliana* distal root stem cell fate regulation via dynamic differential subnuclear localization of transcription factor complexes integrating environmental cues through ELF3. **a:** In response to environmental cues such as light, daylength, or temperature, ELF3 (blue) redistributes into nuclear condensates. This process is facilitated and stabilized by PIFs (yellow), which promote ELF3 nuclear accumulation. **b:** In the root stem cell niche, PLT3 (green) and WOX5 (red) are constitutively present in the nucleus, where PLT3 forms PrD-dependent condensates. Upon environmental stimulation, ELF3 and PIFs co-localize with PLT3 in nuclear condensates in specific SCN cell types. PLT3 recruits WOX5 into these condensates. Through dynamic assembly and disassembly of these PrD-driven condensates, environmental information is integrated at the protein level to modulate QC activity and maintain root SCN homeostasis.

## Materials and methods

### Plant constructs cloning

The GreenGate cloning method (Lampropoulos et al. 2013) was used to create the expression vectors in Supplementary table 4. The sequences upstream of the ATG start codon of ELF3 (3542) and PLT3 (4494) were used as promoter regions. The promoters, as well as the Cytomegalovirus (CMV) promoter, a strong constitutive promoter for expression in mammalian cells, were cloned in the GreenGate entryvector pGGA000 via BsaI restriction and ligation. The GreenGate promoter module carrying the β-estradiol inducible cassette was provided (Denninger et al. 2019). Internal BsaI restriction sites in the PIF4 sequence were removed by PCR amplification using primers with an altered nucleotide sequence at this site (Supplementary Table 1), resulting in two gene fragments that were subsequently reconnected by an overlap extension PCR. The CDS of ELF3, ELF3ΔPrD, PLT3, PLT3ΔPrD, PIF3 and PIF4 were cloned in entry vector pGGC000 and the FPs mVenus and mCherry in pGGD000 via BsaI restriction and ligation. The C-tag module carrying the FP Cerulean (pBLAD002) was a kind gift from Dr. Andrea Bleckmann, University of Regensburg. All entry vectors were confirmed by sequencing. The expression cassettes were created with a GreenGate reaction using pGGZ001 as destination vector. The correct assembly of the modules was confirmed by sequencing. All primer used for GreenGate cloning can be found in Supplementary Table 1.

The Gateway® cloning method (Invitrogen™, Thermo Fisher Scientific Inc.) was used to create the expression vectors in Supplementary table 5. The CDS of ELF3, ELF3ΔQ, PLT3, PLT3ΔQ and mCherry were cloned into pENTR/D-TOPO®. The entry vectors were confirmed by sequencing. The Gateway destination vector carrying the mVenus (pRD04) originates from pMDC7 (Curtis and Grossniklaus 2003) that contains the cassette for β-estradiol inducible expression in planta. The mVenus was introduced via restriction/ligation C-terminally to the Gateway cloning site. The destination vectors with C-terminally located GFP (pABindGFP) and mCherry (pABindmCherry) were described before (Bleckmann et al. 2010). The Gateway destination vectors for mammalian cell culture (pH-Ch-N and pH-mRuby2-N) were a kind gift from Christian Hoischen, Leibniz Institute on Aging -Fritz-Lipmann-Institute, Jena. The mammalian Gateway destination vector carrying the mVenus for C-terminal fusion (pRD10) is based on these vectors and the mVenus was introduced via restriction/ligation. A subsequent LR-reaction of entry and destination vectors was carried out to create the expression vectors. The Gateway expression vectors were verified by test digestion. All primer used for Gateway cloning can be found in Supplementary Table 2.

The domain deletion variant of ELF3 (ELF3ΔPrD) was created by amplifying the sequences upstream and downstream of the deletion. The PrDs were replaced by a 27 aa linker (aagaaggagggaaaaaggagaaaaaga). Fragments and linker were reconnected by overlap extension PCR. The PLT3 deletion variant PLT3ΔPrD has been described before (Burkart et al. 2022).

### Plant work

All *A. thaliana* lines used in this study were in the Columbia (Col-0) background. The *elf3-1* and *plt3-1* single mutants have been previously described (Zagotta et al. 1992; Galinha et al. 2007). The homozygous *elf3-1*, *plt3-1* double mutant was created by crossing (Supplementary table 7). The homozygous *plt3-1* genotype was confirmed by PCR (primer pairs see Supplementary table 3), to obtain the homozygous *elf3-1* genotype, a selection regarding the elongated hypocotyl phenotype was performed. All transgenic lines in this study were created by floral dip as described previously (Zhang et al. 2006). Homozygous lines were confirmed by hygromycin selection. The transgenic line expressing pPLT3::PLT3-mV in the *plt3-1* mutant background has been described previously (Burkart et al. 2022). Plants for crossing, floral dips, genotyping, and seed amplification were grown on soil in phytochambers under constant light or long day (16 h light/ 8 h dark) conditions at 21 °C. For microscopy, a fume-sterilization (50 ml 13 % sodium hypochlorite (v/v) + 1 ml hydrochloric acid) of the Arabidopsis seeds was done beforehand. Sterile seeds were then imbedded in 0.2 % (w/v) agarose, stratified for 2 days at 4 °C and plated on GM agar plates (1/2 strength Murashige Skoog medium including Gamborg B5 vitamins, 1.2 % (w/v) plant agar, 1 % (w/v) sucrose, supplemented with 0.05 % (w/v) MES hydrate). Afterwards, seedlings were grown under continuous light at 21 °C for 5 days. For root imaging, cell walls were stained with either propidium iodide or FM4-64 dye (Invitrogen™, Thermo Fisher Scientific Inc.) as described previously (Burkart et al. 2022).

### Hypocotyl length measurements

Seeds were sterilized and plated on GM agar plates like described above (see Plant work). Afterwards, seedlings were grown in short day (8 h light/ 16 h dark) conditions at 21 °C for seven days. Photos were taken and hypocotyl length was measured with Fiji (Schindelin et al. 2012). Box plots were created with R (version 4.4.1).

### Quantification of leaf area and leaf number

Six seedlings of each genotype were sterilized and plated on GM agar plates and grown in short day (8 h light/ 16 h dark) conditions at 21 °C for seven days. They were then transferred on soil and continued growing in short day (8 h light/ 16 h dark) conditions at 21 °C for sixteen days. Afterwards the phytochamber was switched to long day (16 h light/ 8 h dark, 21 °C) conditions. Pictures were taken from 30 days old plants. The leaves were counted and the leaf area was measured with Fiji (Schindelin et al. 2012). Box plots were created with R (version 4.4.1).

### N. benthamiana infiltration

For transient gene expression, leaves of *N. benthamiana* plants were infiltrated with an *Agrobacterium tumefaciens* strain as described previously (Burkart et al. 2022).

### SCN staining

The combined mPS-PI-and EdU-staining was performed like described before (Burkart et al. 2022). The 2D plots were created with Origin 2020 (OriginLab Corporation).

### Cell culture

All work was done under sterile conditions. Human epithelial cells, HEp-2 (epithelial larynx carcinoma, ATCC-CCL-23) were routinely cultured in Dulbeccós modified Eaglés medium (DMEM, Thermo Fisher Scientific Inc.) at 37 °C under 5 % CO2. Medium was supplemented with 10 % foetal calf serum (FCS), 100 U/ml penicillin and 100 µg/ml streptomycin. For splitting, cells were washed with 5 ml DPBS-Buffer (_’_Dulbecco’s Phosphate Buffered Saline‘) and subsequently incubated for 5 min in 2 ml Trypsin-solution (0.25 % Trypsin, 0.02 % EDTA in DPBS-Buffer) in an incubator (37 °C, 5 % CO2). Trypsinisation was stopped by adding 2 ml medium and 1 ml of the cell-suspension was transferred in a new flask with 7 ml medium. For microscopy, cells were transfected in 2 ml-imaging dishes with the FuGENE® HD transfection reagent (Promega Corporation) according to the manufacturers protocol and a transfection reagent to DNA ratio of 2:1.

### Microscopy

Imaging was carried out with a ZEISS LSM780, LSM880, or LSM980. Fluorescent dyes were excited, and emission was detected as follows: Cerulean was excited at 458 nm and emission was detected at 460-510 nm. mVenus was excited at 514 nm and emission was detected at 517-560 nm, or for co-expression with red dyes excited at 488 nm and detected at 500-560 nm. Alexa Fluor® 488 was excited at 488 nm and emission was detected at 490-560 nm. PI was excited at 561 nm and emission was detected at 590-710 nm. FM4-64 was excited at 514 nm or 561 nm and emission was detected at 670-760 nm. mCherry and mRuby2 were excited at 561 nm and emission was detected at 590-640 nm. To avoid cross talk, imaging of more than one fluorophore was carried out in the sequential mode.

### Intensity measurements

The intensity measurements were performed in a transient *N. benthamiana* experiments. To analyze the ratio of the nuclear and cytoplasmic fraction of the ELF3-mVenus variants and the ELF3-mVenus co-expressed PLT3-mCherry variants, images of nuclei and cytoplasm of the same cell were acquired with constant settings for the mVenus channel at the ZEISS LSM880. Fiji (Schindelin et al. 2012) was used for data analysis, and mean intensities of ROIs were measured in the nucleus and corresponding cytoplasm. The intensity ratio was plotted in a box plot created with R (version 4.4.1).

### Body tracking

3D time series of the nuclear PLT3-mVenus condensates and the cytoplasmic ELF3-mVenus condensates were acquired with the ZEISS LSM880 airyscan fast mode in transient N. benthamiana experiments. The tracking and data analysis were done with Imaris (version 9.1.2, Bitplane, Oxford Instruments plc). For the PLT3-mVenus nuclei, a shift correction was performed beforehand with Imaris (version 9.1.2, Bitplane, Oxford Instruments plc).

### FLIM measurements

The FLIM measurements in *N. benthamiana* leaf epidermal cells were performed as described previously (Burkart et al. 2022). The fluorophores mVenus and mCherry were used as donor-acceptor pair in the interaction experiments. The mVenus donor was excited with a linearly polarized diode laser (LDH-D-C-485) at 485 nm and a pulse frequency of 32 MHz. The excitation power was adjusted to 0.5-1 µW at the objective (C-Apochromat 40×/1.2 W Corr M27, ZEISS). FLIM data analysis was done as described previously (Burkart et al. 2022). Subsequently the intensity weighted lifetime was has been taken for further analysis. Box and scatter plots were created with R (version 4.4.1).

### Prediction of protein domains

The PrDs in ELF3 and PLT3 were predicted using the PLAAC application (Lancaster et al. 2014). The nuclear localization signals (NLS) in ELF3 and PLT3 were predicted with cNLS Mapper (Kosugi et al. 2009).

### Cell-free expression and immunoprecipitation

ELF3 (AT2G25930.1, Uniprot ID: O82804), PLT3 (AT5G10510.6, Uniprot ID: F4KGW9), PIF3 (AT1G09530.6, Uniprot ID: O80536) and PIF4 (AT2G43010.1, Uniprot ID: Q8W2F3) full length were PCR-amplified and inserted into the pTnT (Promega L5610) vectors pTnT-C-3XFLAG, pTnT-N-7XMyc, pTnT-N-StrepTagII and pTnT-N-V5. PLT3 has a mutation at position 284 from a glycine (in F4KGW9) to an arginine.

The vectors were used for *in vitro* protein production using the TnT SP6 High-Yield Wheat Germ Protein Expression System (Promega, catalogue No. L3260) according to the manufacturer’s instructions. In brief, 1 µg of each indicated plasmid was mixed with 20 µl TnT SP6 High-Yield Wheat Germ Master Mix and completed to 35 µl with nuclease-free water. The reaction was incubated at 25°C for 2 h. 5 µl of each reaction was used as the input for Western blots. The remaining reaction was topped up with 100 µl pulldown buffer (10 mM Tris pH 7.5, 300 mM NaCl, 1% Triton X-100, 1.5 mg/ml BSA) and 10 µl anti-FLAG magnetic beads (Anti-FLAG M2 magnetic beads; catalogue No. M8823, Merck Millipore) and incubated 1 h at 4°C. The anti-FLAG magnetic beads were washed for 5 x 2 min under agitation with 500 µl pulldown buffer. Finally, the beads were resuspended in 20 µl phosphate-buffered saline and 10 µl SDS–PAGE loading dye and boiled for 5 min at 95°C and separated on a 12%SDS PAGE, which was used for Western blots using anti-FLAG [monoclonal ANTI-FLAG M2-peroxidase (HRP) antibody; catalogue No. A8592, Sigma-Aldrich; 1:10 000 dilution], anti-Myc (Myc Tag Monoclonal Antibody, HRP; catalogue No. R95125, Invitrogen; 1:5000 dilution), anti-StrepTagII (StrepTag II Antibody HRP Conjugate; catalogue No. 71591-M, Sigma-Aldrich; 1:5000 dilution) and anti-V5 (V5 Tag Monoclonal Antibody, HRP; catalogue No. R961-25, Invitrogen; 1:10 000 dilution) antibodies.

### Cloning and expression of recombinant proteins

Soluble PLT3 PrD2 (PLT3 residues 415-569) and ELF3 PrD constructs (Q7; residues 388– 625; AT2G25930, *Arabidopsis thaliana* Col-0) were identified using the Esprit library (Yumerefendi et al. 2010; An et al. 2011a; An et al. 2011b; Hart and Waldo 2013; Mas and Hart 2017). All constructs were expressed in *Escherichia coli* BL21-CodonPlus-RIL cells (Agilent) at 18 °C, induced with 1 mM IPTG for 16 h. Cells were resuspended in 100 mM Bis-Tris-propane pH 9.4, 300 mM NaCl, 20 mM imidazole, 1 mM TCEP, and EDTA-free protease inhibitors (ThermoFisher). The supernatant was loaded onto a 1 mL Ni-NTA affinity column equilibrated in lysis buffer (without protease inhibitors), washed with the same buffer and subsequently with a high-salt buffer (100 mM Bis-Tris-propane pH 9.4, 1 M NaCl, 20 mM imidazole, 1 mM TCEP). Bound proteins were eluted using 100 mM Bis-Tris-propane pH 9.4, 300 mM NaCl, 300 mM imidazole, and 1 mM TCEP. Purity was confirmed by SDS–PAGE, and fractions containing the target protein were pooled and dialyzed for ∼2 h at 4 °C against 50 mM Bis-Tris-propane pH 9.4, 500 mM NaCl, and 1 mM TCEP. Final protein concentrations ranged from 3 to 8 mg mL⁻¹. Bis-Tris-propane was chosen as the buffering agent due to its broad effective pH range.

### SAXS

SAXS measurements were carried out at the European Synchrotron Radiation Facility (ESRF) on the BioSAXS beamline BM29 (69, 70). Protein solutions were prepared at 4 mg mL⁻¹ for ELF3 PrD Q7 and 3.3 mg/mL PLT3 PrD2 and run on a S200 Increase 3.2 300. Data were collected using a Pilatus 2M detector (Dectris) at a sample–detector distance of 2.81 m. Initial processing was performed with the Dahu pipeline (72, 73), producing automatically normalized, radially integrated 1-D scattering profiles for each frame. Further processing was performed in Scatter IV (74).

### MALLS

50 µl of PLT3 at 7mg/mL were loaded onto an S200 Increase size-exclusion column (Superdex 200 Increase10/300 GL, GE Healthcare) at a flow rate of 0.5 mL min−1. The column was preequilibrated with 50 mM Bis Tris propane at pH 9.4, 1 M NaCl, 1 mM TCEP and connected to a Hitachi Elite LaChrom UV detector and LAChrome Pump L-2130, a multiangle laser light-scattering detector (DAWN HELEOS II, Wyatt Technology Corporation) and a refractive-index detector (Optilab T-rEX, Wyatt Technology Corporation). The data were processed with the ASTRA 6.1.7.17 software (Wyatt Technology Corporation).

## Acknowledgements

We thank Philipp Wuthenow and Milena Twarz for supporting parts of the lab work. We would like to acknowledge funding by the Deutsche Forschungsgemeinschaft (DFG) to Y.S. via grant 490818643 (STA 1212/6-1) and the DFG FUGG program (project 553896058, Fluorescence Imaging Microscopy Microscope). We would like to acknowledge the Center for Advanced Imaging (CAI) at Heinrich Heine University for support with imaging.

This project received support from the ANR (ANR-19-CE20-0021, ANR-21-CE11-0037 and ANR-25-CE20-6732) and GRAL, a program from the Chemistry and Biology Health Graduate School of the University Grenoble Alpes (ANR-17-EURE-0003). We thank the microscopy facility MuLife of IRIG/DBSCI, funded by CEA Nanobio and GRAL LabEX (ANR-10-LABX-49-01) financed within the University Grenoble Alpes graduate school CBH-EUR-GS (ANR-17-EURE-0003). The SAXS dilute and condensed phase experiments were performed on BM29 at the European Synchrotron Radiation Facility (ESRF), Grenoble, France. Financial support was provided by Instruct-ERIC (PID 13317). This work used the platforms of the Grenoble Instruct-ERIC center (ISBG; UAR 3518 CNRS-CEA-UGA-EMBL) within the Grenoble Partnership for Structural Biology (PSB), supported by FRISBI (ANR-10-INBS-0005-02). The CBS is a member of the France-BioImaging (FBI), national infrastructure supported by the French National Research Agency (ANR-10-INBS-04-01) and of the GIS IBISA (Infrastructures en Biologie Santé et Agronomie).

L.C. received support from the French National Research Agency (ANR) (ANR-21-CE11-0037), and from the CNRS Momentum program (2017). The CBS is a member of France-BioImaging (FBI), a national infrastructure supported by the ANR (ANR-10-INBS-04-01), and of the GIS IBiSA (Infrastructures en Biologie Santé et Agronomie).

## Notes

### Competing Interest Statement

The authors have declared no competing interest.

### Summary of Updates

1.The author list and affiliations have been updated. 2.The abstract has been updated. 3.The main figures have been streamlined from 12 to 10. The in vitro biophysical characterization data (former Figures 7 and 8) have been reassessed, condensed, and moved to Supplementary Figure 3, and figure numbering has been updated accordingly throughout the manuscript. 4.A new pull-down experiment has been added as Supplementary Figure 4, providing additional support for the ELF3-PLT3-PIF3-PIF4 interactions. 5.Supplemental files updated in accordance with the revised figures and tables.

