## supplemental information for "EARLY FLOWERING 3 (ELF3): a novel role in integrating environmental stimuli with root stem cell niche maintenance"

### Supplementary figures and tables

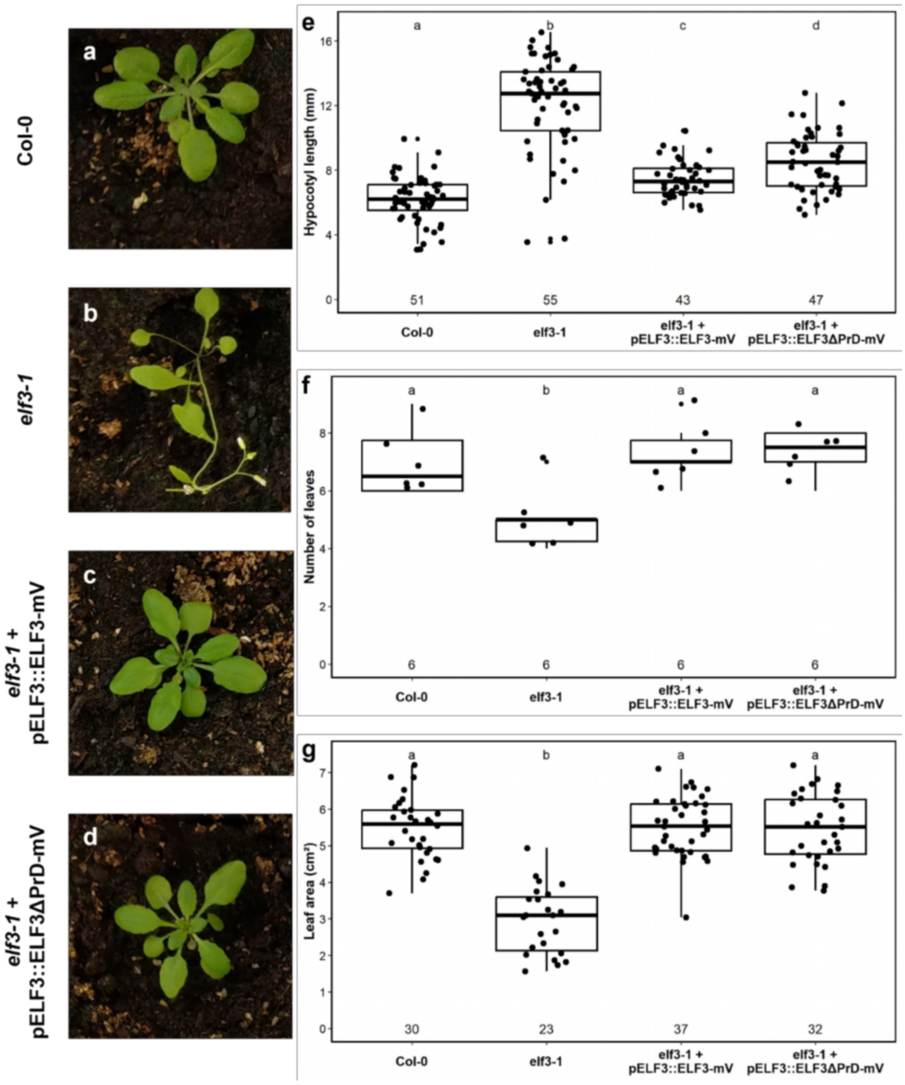


**Supplementary figure 1: Phenotypes of Col-0 wild type, *elf3-1* mutant and transgenic rescue lines**

**a-d:** Representative images of 30-day old plants of Col-0 wild type (a), *elf3-1* mutant (b), transgenic elf3 + pELF3::ELF3-mV line (c) and elf3 + pELF3::ELF3ΔPrD-mV line, expressing the PrD deletion variant (d), grown in short day conditions. **e-g:** Scatter and box plots presenting the hypocotyl lengths (mm) of 7-day old seedlings of the indicated genotypes **(e)**, number of leaves **(f)**, and leaf area ($cm^2$) **(g)** of 30-day old plants. Each dot represents a single measurement, boxes indicate the 25–75% percentile, whiskers show 1.5 x the interquartile range, the line marks the median, and the square denotes the mean. Replicate numbers are shown below each group. Statistical analyses were performed using one-way ANOVA followed by Tukey's HSD post hoc test for normally distributed data, or Kruskal–Wallis test followed by Dunn's post hoc test with multiple testing correction when normality assumptions were not met. Groups sharing the same letter are not significantly different ($\alpha = 0.01$).

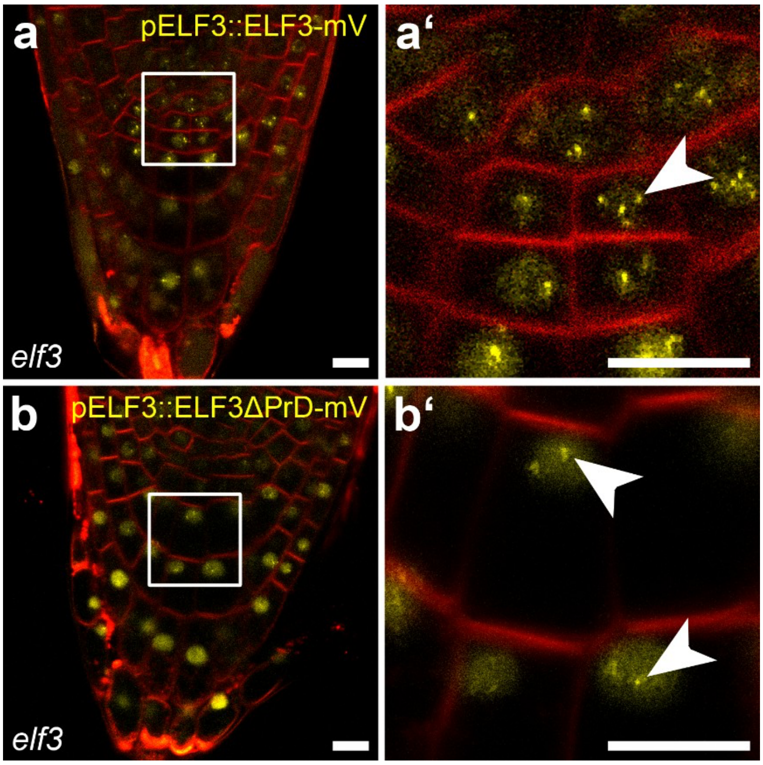


**Supplementary figure 2: Localization of ELF3-mV and ELF3ΔPrD-mV in the RAM of *A. thaliana***

**a-a',** Localization of endogenously expressed ELF3-mV (yellow) in the *elf3* mutant root. Cell walls are stained with FM4-64 (red). **a'** is a magnification of **a**, indicated by the white square in **a**. **b-b',** Localisation of an endogenously expressed ELF3-PrD deletion variant (ELF3ΔPrD-mV) in the *elf3* mutant root. Cell walls are stained with FM4-64 (red). **b'** is a magnification of **b**, indicated by the white square in **b**. White arrowheads point at NBs. Scale bars represent 5 μm. mV = mVenus; PrD = prion-like domain.

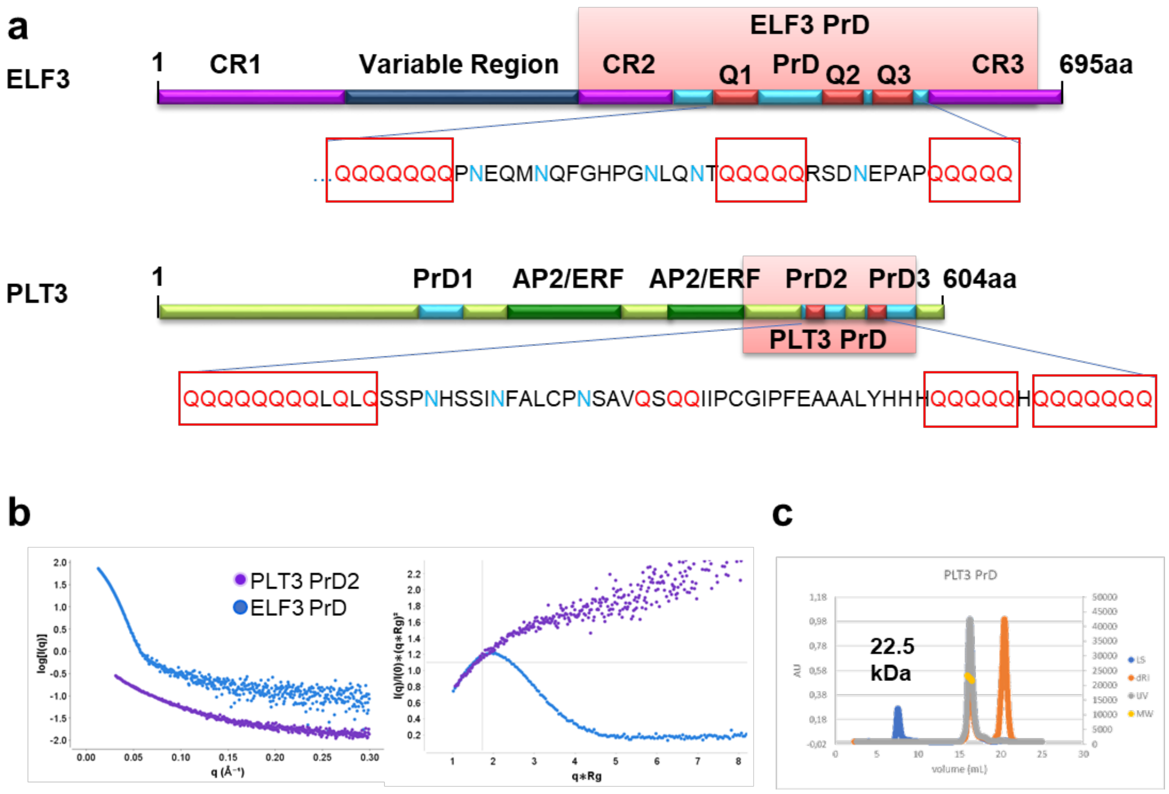


**Supplementary figure 3: ELF3 and PLT3 prion-like domains interact in vitro and colocalize to condensates.**

**a:** Schematic of the ELF3 (top) and PLT3 (bottom) primary amino acid sequence with domains and regions indicated. The polyQregions are shown as a zoom with glutamines highlighted in red. Conserved Regions (CR), Variable Region, Prion-like Domain (PrD), polyQ (Q), APETALA2/Ethylene Response Facto (AP2/ERF). **b:** Left, overlay of X-ray scattering curves for PLT3 PrD(purple), ELF3 PrD (blue) and a mixture of PLT3/ELF3 PrDs(brown). Right, Kratky plot for PLT3 PrD, ELF3 PrD and a mixture of PLT3/ELF3 PrDs colored as per left. The plateau for PLT3 PrDindicates a flexible unfolded protein. The bell shaped curves for ELF3 PrD and the mixture indicate a globular species in solution. **c:** Size-exclusion multi-angle laser light scattering chromatogram for PLT3 indicating a monodisperse and monomeric species of 22.5 kDa in solution. LS-light scattering, dRI-differential refractive index, UV-ultraviolet 280 nM, MW-molecular weight

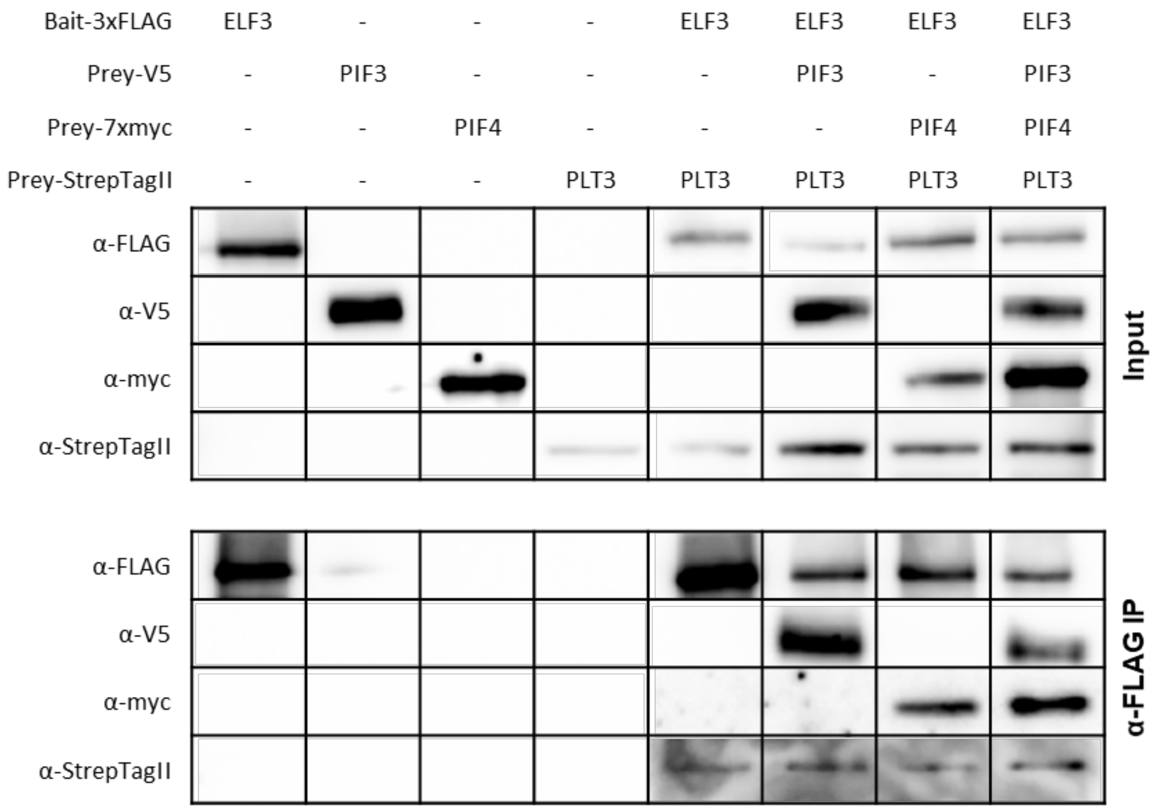


**Supplementary figure 4: Pull-down experiment demonstrating interaction between ELF3, PLT3, PIF3 and PIF4.**

All proteins were expressed using in vitro transcription translation and different N- or C-terminal tags as indicated on top. The upper table shows the input and the lower the anti-FLAG IP.

### Supplementary table 1: List of primers for GreenGate cloning

| gene ID | alias | primer name | orientation | sequence |
|---|---|---|---|---|
| **Promoter modules** | | | | |
| AT2G25930 | pELF3 | PW_GG_p-ELF3 F | F | TTTGGTCTCAACCTTTTCGATCAAAGCAGCAGATTC |
| | | PW_GG_p-ELF3 R | R | AAAGGTCTCATGTTCACTCACAATTCACAACC |
| AT5G10510 | pPLT3 | RD_GG pPLT3 F V2 | F | AAAGGTCTCAACCTAATTTTAACGTATTCTTTC |
| | | RD_GG pPLT3 R V2 | R | AAAGGTCTCATGTTAAACTTTCTTATAAAAACAATT |
| - | pCMV for *HEp-2* | PW_GG_P-CMV F | F | AAAGGTCTCAACCTCCGCCATGCATTAGTTATTAATAG |
| | | PW_GG_P-CMV R | R | AAAGGTCTCATGTTGATCTGACGGTTCACTAAACC |
| **CDS modules** | | | | |
| AT2G25930 | ELF3 | RD_GG ELF3 F | F | AAAGGTCTCAGGCTTAATGAAGAGAGGGAAAG |
| | | RD_GG ELF3 R | R | AAAGGTCTCACTGAAGGCTTAGAGGAGTCATAG |
| | ELF3ΔPrD | PW_GG_p-ELF3 F | F | TTTGGTCTCAACCTTTTCGATCAAAGCAGCAGATTC |
| | | RD_ELF3deltaPrD1 R | R | CCAGCTGCCAACCTCCCAACTAC |
| | | RD_ELF3 linker F | F | GAGGTTGGCAGCTGGTGCTGC |
| | | RD_ELF3 linker R | R | CGCGGTGCTCCTGCGG |
| | | RD_ELF3deltaPrD2 F | F | AGGAGCACCGCGAGCAAGAAAG |
| | | PW_GG_p-ELF3 R | R | AAAGGTCTCATGTTCACTCACAATTCACAACC |
| AT5G10510 | PLT3 | RD_GreenGate PLT3 F | F | AAAGGTCTCAGGCTTAATGGAGATGTTGAG |
| | | RD_GreenGate PLT3 R | R | AAAGGTCTCACTGAGTAAGACTGATTAGGC |
| | PLT3ΔPrD | RD_GreenGate PLT3 F | F | AAAGGTCTCAGGCTTAATGGAGATGTTGAG |
| | | RD_PLT3ΔPrD1 CDS1 R | R | CCAGCTGCAACACCAAGTGACAAAG |
| | | RD_PLT3ΔPrD1 linker F | F | CTTGGTGTTGCAGCTGGTGCTG |
| | | RD_PLT3ΔPrD1 linker R | R | GTCTTCTCTGCTCCTGCGGCAG |
| | | RD_PLT3ΔPrD1 CDS2 F | F | CGCAGGAGCAGAGAAGACAGATTCTG |
| | | RD_GG PLT3ΔPrDs R | R | AAAGGTCTCACTGAGTGAAGTTGATGATGAC |
| AT1G09530 | PIF3 | RD_GG PIF3 CDS F | F | AAAGGTCTCAGGCTTAATGCCTCTGTTTGAG |
| | | RD_GG PIF3 CDS R | R | AAAGGTCTCACTGACGACGATCCACAAA |
| AT2G43010 | PIF4 | RD_GG PIF4 CDS F | F | AAAGGTCTCAGGCTTAATGGAACACCAAG |
| | | RD_GG PIF4 CDS R | R | AAAGGTCTCACTGAGTGGTCCAAACGAG |
| | | RD_GG PIF4 Mitte F | F | GATCCCCTCCAAAGACCAACCTC |
| | | RD_GG PIF4 Mitte R | R | GAGGTTGGTCTTTGGAGGGGATC |
| **C-tag modules** | | | | |
| - | mVenus | RD_GreenGate mVenus C-tag F | F | AAAGGTCTCATCAGCAATGGTGAGCAAGG |
| | | RD_GreenGate mVenus C-tag R | R | AAAGGTCTCAGCAGTTACTTGTACAGCTC |
| - | mCherry | RD_GG mCherry C-tag F | F | AAAGGTCTCATCAGCAATGGTGAGCAAGG |
| | | RD_GG mCherry C-tag R | R | AAAGGTCTCAGCAGTTACTTGTACAGCTCGTC |
| **Terminator modules** | | | | |

| | | | | |
|---|---|---|---|---|
| - | SV40-polyA for *HEp-2* | PW_GG_SV40pA F | F | AAAGGTCTCACTGCGCCATACCACATTTGTAGAG |
| | | PW_GG_SV40pA R | R | AAAGGTCTCATAGTCGCCTTAAGATACATTGATGAG |
| **Plant resistance modules** | | | | |
| - | SV40 ori for *HEp-2* | PW_GG_SV40o F | F | AAAGGTCTCAACTAGGTGTGGAAAGTCCCC |
| | | PW_GG_SV40o R | R | AAAGGTCTCAATACGGCCTCCAAAAAAGCC |

### Supplementary table 2: List of primers for Gateway Cloning

| gene ID | alias | primer name | orientation | sequence |
|---|---|---|---|---|
| AT2G25930 | ELF3 | PW ELF3 CACC F | F | CACCATGAAGAGAGGGAAAGATGA |
| | | ELF3 ohne Stopp R | R | AGGCTTAGAGGAGTCATAGC |
| | ELF3 for *HEp-2* | PW ELF3 CACC F | F | CACCATGAAGAGAGGGAAAGATGA |
| | | ELF3 R-stop +2BP | R | CCAGGCTTAGAGGAGTCATAGC |
| AT5G10510 | PLT3 | YS_PLT3 F CACC | F | CACCATGGAGATGTTGAGGTCATCTGATCAGTCTCA |
| | | YS_PLT3 R -stop | R | GTAAGACTGATTAGGCCAGAGGAAG |
| | PLT3ΔQ | YS_PLT3 F CACC | F | CACCATGGAGATGTTGAGGTCATCTGATCAGTCTCA |
| | | RD_PLT3 Seg1 R | R | GAGATGAGAAATGGTGAAGTTGATGATGAC |
| | | RD_PLT3 Seg2 F | F | CTTCACCATTTCTCATCTCCTAATCACAGTAGC |
| | | RD_PLT3 Seg2 R | R | GAAGAAGTTGTGGTGGTGGTAAAGAGCAG |
| | | RD_PLT3 Seg3 F | F | CACCACCACAACTTCTTCCAGCATTTTCC |
| | | YS_PLT3 R -stop | R | GTAAGACTGATTAGGCCAGAGGAAG |
| | PLT3 for *Hep-2* | YS_PLT3 F CACC | F | CACCATGGAGATGTTGAGGTCATCTGATCAGTCTCA |
| | | RD_PLT3 R -stop +2bp | R | AAGTAAGACTGATTAGGCCAGAGGAAG |
| - | mVenus | RD_mVenus PacI F | F | CCCTTAATTAAAATGGTGAGCAAGGGCGAGG |
| | | MAS-Turquoise-SpeI-R | R | TTTACTAGTTACTTGTACAGCTCGTCCATGC |
| - | mVenus for *Hep-2* | YS_mVenus AgeI F | F | AAACCGGTATGGTGAGCAAGGGCGAGGA |
| | | YS_mVenus MfeI R | R | AAACAATTGTTACTTGTACAGCTCGTCCATGCCG |

### Supplementary table 3: List of genotyping primers

| gene ID | alias | primer name | orien-tation | sequence |
|---|---|---|---|---|
| AT5G10510 | *plt3-1* | GK_PLT3L | F | TTGTGATTTGCCATTGACTAAAGGT |
| | | GK_PLT3R | R | GAAAACAGTCCAATGGTCTCACATC |
| | | SALK LBB1V2 | | AAACCAGCGTGGACCGCTTGCTGCAACTCT |

### Supplementary table 4: List of GreenGate expression vectors used in this study

| construct | promoter | N-tag | CDS | C-tag | term-inator | plant sel. marker | destination vector | bacterial sel. marker | plasmid ID |
|---|---|---|---|---|---|---|---|---|---|
| | module A | module B | module C | module D | module E | module F | module Z | | |
| **pELF3::ELF3-mV** | ELF3 promoter | Ω-element (pGGB002) | ELF3 | mVenus | tUBQ10 (pGGE009) | Hyg (pGGF005) | pGGZ001 | Spec | pRD99 |
| **pELF3::ELF3ΔPrD-mV** | ELF3 promoter | Ω-element (pGGB002) | ELF3 ΔPrD | mVenus | tUBQ10 (pGGE009) | Hyg (pGGF005) | pGGZ001 | Spec | pRD124 |
| **35S::ELF3-mV** | 35S (pGGA004) | Ω-element (pGGB002) | ELF3 | mVenus | tUBQ10 (pGGE009) | Hyg (pGGF005) | pGGZ001 | Spec | pRD98 |
| **inducible ELF3ΔPrD-mV** | Ubi-XVE oLexA-35S | Ω-element (pGGB002) | ELF3 ΔPrD | mVenus | tUBQ10 (pGGE009) | Hyg (pGGF005) | pGGZ001 | Spec | pRD119 |
| **pPLT3:: PLT3-mV** | PLT3 promoter | Ω-element (pGGB002) | PLT3 | mVenus | tUBQ10 (pGGE009) | Hyg (pGGF005) | pGGZ001 | Spec | pRD73 |
| **pPLT3:: PLT3ΔPrD-mV** | PLT3 promoter | Ω-element (pGGB002) | PLT3 ΔPrD | mVenus | tUBQ10 (pGGE009) | Hyg (pGGF005) | pGGZ001 | Spec | pRD125 |
| **inducible PLT3-C** | Ubi-XVE oLexA-35S | Ω-element (pGGB002) | PLT3 | Cerulean | tUBQ10 (pGGE009) | Hyg (pGGF005) | pGGZ001 | Spec | pRD80 |
| **inducible PLT3ΔPrD-mV** | Ubi-XVE oLexA-35S | Ω-element (pGGB002) | PLT3 ΔPrD | mVenus | tUBQ10 (pGGE009) | Hyg (pGGF005) | pGGZ001 | Spec | pRD106 |
| **inducible PLT3ΔPrD-mC** | Ubi-XVE oLexA-35S | Ω-element (pGGB002) | PLT3 ΔPrD | mCherry | tUBQ10 (pGGE009) | Hyg (pGGF005) | pGGZ001 | Spec | pRD138 |
| **inducible PIF3-mC** | Ubi-XVE oLexA-35S | Ω-element (pGGB002) | PIF3 | mCherry | tUBQ10 (pGGE009) | Hyg (pGGF005) | pGGZ001 | Spec | pLC08 |
| **inducible PIF3-mV for HEp-2** | pCMV | Ω-element (pGGB002) | PIF3 | mVenus | SV40-polyA | SV40 ori | pGGZ001 | Spec | pRD93 |
| **inducible PIF3-C for HEp-2** | pCMV | Ω-element (pGGB002) | PIF3 | Cerulean | SV40-polyA | SV40 ori | pGGZ001 | Spec | pRD94 |
| **inducible PIF4-mV** | Ubi-XVE oLexA-35S | Ω-element (pGGB002) | PIF4 | mVenus | tUBQ10 (pGGE009) | Hyg (pGGF005) | pGGZ001 | Spec | pLC10 |
| **inducible PIF4-mC** | Ubi-XVE oLexA-35S | Ω-element (pGGB002) | PIF4 | mCherry | tUBQ10 (pGGE009) | Hyg (pGGF005) | pGGZ001 | Spec | pLC11 |
| **inducible PIF4-mV for HEp-2** | pCMV | Ω-element (pGGB002) | PIF4 | mVenus | SV40-polyA | SV40 ori | pGGZ001 | Spec | pRD95 |
| **inducible PIF4-C for HEp-2** | pCMV | Ω-element (pGGB002) | PIF4 | Cerulean | SV40-polyA | SV40 ori | pGGZ001 | Spec | pRD96 |

**Supplementary table 5: List of Gateway expression vectors used in this study**

| construct | promoter | CDS | C-tag | termi-nator | plant sel. marker | destination vector | bacterial sel. marker | plasmid ID |
|---|---|---|---|---|---|---|---|---|
| **inducible ELF3-mV** | Ubi-XVE oLexA-35S | ELF3 | mVenus | T3A | Hyg | pRD04 | Spec | pPW06 |
| **inducible ELF3ΔQ-mV** | Ubi-XVE oLexA-35S | ELF3ΔQ | mVenus | T3A | Hyg | pRD04 | Spec | pPW47 |
| **inducible ELF3-mV for HEp-2** | Ubi-XVE oLexA-35S | ELF3 (+2bp) | mVenus | T3A | Hyg | pRD10 | Amp | pRD49 |
| **inducible PLT3-mV** | Ubi-XVE oLexA-35S | PLT3 | mVenus | T3A | Hyg | pRD04 | Spec | pRD25 |
| **inducible PLT3-GFP** | Ubi-XVE oLexA-35S | PLT3 | GFP | T3A | Hyg | pABind GFP | Spec | pFB05 |
| **inducible PLT3-mC** | Ubi-XVE oLexA-35S | PLT3 | mCherry | T3A | Hyg | pABind mCherry | Spec | pFB06 |
| **inducible PLT3ΔQ-mV** | Ubi-XVE oLexA-35S | PLT3ΔQ | mVenus | T3A | Hyg | pRD04 | Spec | pRD57 |
| **inducible PLT3ΔQ-mC** | Ubi-XVE oLexA-35S | PLT3ΔQ | mCherry | T3A | Hyg | pABind mCherry | Spec | pRD81 |
| **inducible PLT3-mV for HEp-2** | Ubi-XVE oLexA-35S | PLT3 (+2bp) | mVenus | T3A | Hyg | pRD10 | Amp | pRD11 |
| **inducible PLT3-mRb2 for HEp-2** | Ubi-XVE oLexA-35S | PLT3 (+2bp) | mRuby2 | T3A | Hyg | pH-mRuby2-N | Amp | pRD13 |
| **inducible PLT3-mC for HEp-2** | Ubi-XVE oLexA-35S | PLT3 (+2bp) | mCherry | T3A | Hyg | pH-Ch-N | Amp | pRD36 |
| **inducible free mCherry** | Ubi-XVE oLexA-35S | mCherry | - | T3A | Hyg | pMDC7 | Spec | - |

**Supplementary table 6: List of Gibbson assembly cloning vectors and primers used in this study**

| gene ID | alias | Plasmid + tag | orientation | sequence |
|---|---|---|---|---|
| AT2G25930.1 | ELF3 | pTNT_C_3xFLAG | F | AGCTACTTGTTCTTTTTGCACTCGAGATGAAGAGAGGGAAAGATG |
| | | | R | GAGCTCCCAGATCCGCTACCGTCGACTCTAGAAGGCTTAGAGGAGTCATAG |
| | | | F | GGTAGCGGATCTGGGAGC |
| | | | R | TGCAAAAAGAACAAGTAGC |
| AT5G10510.6 | PLT3 | pTnT_N_Strep-tag II | F | CACCCGCAGTTCGAAAAAGGAAGCGGCTCAATGGAGATGTTG |

| | | | | |
|---|---|---|---|---|
| | | | R | TTTTTGCGGCCGCCCGGGTCGACTCTAGAGGT AAGACTGATTAGGCCAGAGGAAGAAC |
| | | | F | CTCTAGAGTCGACCCGGGCG |
| | | | R | TGAGCCGCTTCCTTTTTCGAACTG |
| AT1G095 30.1 | PIF3 | pTNT_V5-tag | F | CGATTCTACGGGAAGCGGCTCAATGCCTCTGTT TGAGCTTTTCAGG |
| | | | R | GCTAGTTTTTTCACGACGATCCACAAAACTGAT CAG |
| | | | F | ATCGTCGTGAAAAAACTAGCATAACCCCTTGG GG |
| | | | R | ACAGAGGCATTGAGCCGCTTCCCGTAGAATCG AGACCGAGGAGAG |
| AT2G430 10.1 | PIF4 | pTnT_N_7xMyc-tag | F | AAGAGGACTTGAATGAAATGATGGAACACCAA GGTTGGAGTT |
| | | | R | TTAGAGGCCCCAAGGGGTTAGTGGTCCAAACG AGAACCG |
| | | | F | TAACCCCTTGGGGCCTCTAAAC |
| | | | R | CATTTCATTCAAGTCCTCTTCAGAAATGAGC |

**Supplementary table 7: *Arabidopsis thaliana* mutants and transgenic lines**

| gene ID | alias | reference |
|---|---|---|
| AT2G25930 | *elf3-1* | (Zagotta et al. 1992) |
| AT5G10510 | *plt3-1* | (Galinha et al. 2007) |
| AT2G25930, AT5G10510 | *elf3 plt3* | this study, crossing of *elf3-1* and *plt3-1* |
| AT2G25930 | *pELF3::ELF3-mV (elf3-1)* | this study |
| AT2G25930 | *pELF3::ELF3ΔPrD-mV (elf3-1)* | this study |
| AT5G10510 | *pPLT3::PLT3-mV (plt3-1)* | (Burkart et al. 2022) |

**Supplementary table 8: Spearman's correlation between QC divisions and CSC layers by genotype**

| genotype | average QC cell-divisions per root (± st. dev.) | p-value | number of analysed roots |
|---|---|---|---|
| **Col-0** | -0.19 | 0.01 | 209 |
| ***elf3-1*** | -0.30 | 0.03 | 56 |
| ***plt3-1*** | -0.11 | 0.15 | 172 |
| ***elf3-1, plt3-1*** | -0.28 | 0.05 | 53 |

**Supplementary table 9: Average QC and CSC phenotypes related to** Fehler! Verweisquelle konnte nicht gefunden werden.

| genotype | average QC cell-divisions (± st. dev.) | average CSC layers (± st. dev.) |
|---|---|---|
| **Col-0** | 0.49 ± 0.79 | 0.93 ± 0.58 |
| ***elf3-1*** | 0.55 ± 0.75 | 0.95 ± 0.69 |
| ***plt3-1*** | 0.65 ± 0.77 | 0.82 ± 0.63 |
| ***elf3-1, plt3-1*** | 0.91 ± 0.77 | 0.87 ± 0.70 |

**Supplementary table 10: Average nuclear/cytoplasmic intensity ratio of ELF3-mVenus variant expressing *N. benthamiana* leaf epidermal cells with and without PLT3-mCherry variants.**

| construct | intensity ratio (nucleus/cytoplasm) | standard deviation | number of analysed cells |
|---|---|---|---|
| **ELF3-mVenus** | 2.44 | 1.91 | 12 |
| **ELF3ΔPrD-mVenus** | 3.17 | 2.62 | 10 |
| **ELF3-mVenus + PLT3-mCherry** | 11.86 | 8.54 | 10 |
| **ELF3-mVenus + PLT3ΔPrD-mCherry** | 2.59 | 1.80 | 10 |

**Supplementary table 11: Average hypocotyl length in *elf3-1* rescue experiment**

| genotype | average hypocotyl length | standard deviation | number of analysed seedlings |
|---|---|---|---|
| **Col-0** | 6.11 | 1.36 | 51 |
| ***elf3-1*** | 12.39 | 2.40 | 55 |
| ***elf3-1* + pELF3:ELF3-mV** | 7.38 | 1.01 | 43 |
| ***elf3-1* + pELF3:ELF3ΔPrD-mV** | 8.52 | 1.80 | 47 |

**Supplementary table 12: Average leaf area in *elf3-1* rescue experiments**

| genotype | average leaf area | standard deviation | number of analyzed leaves | number of plants |
|---|---|---|---|---|
| **Col-0** | 5.47 | 0.85 | 30 | 6 |
| ***elf3-1*** | 2.96 | 0.91 | 23 | 6 |
| ***elf3-1* + pELF3:ELF3-mV** | 5.51 | 0.82 | 37 | 6 |
| ***elf3-1* + pELF3:ELF3ΔPrD-mV** | 5.54 | 1.04 | 32 | 6 |

**Supplementary table 13: Average number of leaves in *elf3-1* rescue experiments**

| genotype | average number of leaves | standard deviation | number of plants |
|---|---|---|---|
| **Col-0** | 7.00 | 1.26 | 6 |
| ***elf3-1*** | 5.00 | 1.10 | 6 |
| ***elf3-1* + pELF3:ELF3-mV** | 7.33 | 1.03 | 6 |
| ***elf3-1* + pELF3:ELF3ΔPrD-mV** | 7.33 | 0.82 | 6 |

**Supplementary table 14: Average speed and displacement length of the body-tracking in ELF3-mVenus and PLT3-mVenus expressing *N. benthamiana* epidermal leaf cells**

| construct | mean track speed [µm/s] | variance | track displace-ment length [µm] | variance | number of analyzed condensates | number of analyzed cells |
|---|---|---|---|---|---|---|
| **ELF3-mVenus** | 0.74 | 0.08 | 2.53 | 4.33 | 28493 | 10 |
| **PLT3-mVenus** | 0.06 | 0.001 | 0.50 | 0.16 | 1334 | 9 |

**Supplementary table 15: Average fluorescence lifetimes (τ) and standard deviation (st. dev.) obtained in FLIM experiments**

| donor | acceptor | average $\tau$ [ns] | st. dev. [ns] | Number of analyzed cells |
|---|---|---|---|---|
| **ELF3-mVenus** | - | 2.99 | 0.06 | 60 |
| | **+ free mCherry** | 2.99 | 0.04 | 39 |
| | **+ PLT3-mCherry** | 2.42 | 0.32 | 78 |
| | **+ PLT3ΔPrD-mCherry** | 2.91 | 0.14 | 55 |
| **ELF3ΔPrD-mVenus** | - | 3.03 | 0.04 | 37 |
| | **+ free mCherry** | 3.00 | 0.02 | 36 |
| | **+ PLT3-mCherry** | 2.95 | 0.10 | 54 |
| | **+ PLT3ΔPrD-mCherry** | 2.98 | 0.07 | 35 |

**Supplementary table 16: Average fluorescence lifetimes (τ) and standard deviation (st. dev.) obtained in FLIM experiments with PIF-proteins.**

| donor | acceptor | average $\tau$ [ns] | st. dev. [ns] | Number of analyzed cells |
|---|---|---|---|---|
| PIF3-mVenus | - | 2.87 | 0.08 | 66 |
| | + free mCherry | 2.69 | 0.07 | 33 |
| | + PLT3-mCherry | 2.48 | 0.18 | 39 |
| | + ELF3-mCherry | 1.54 | 0.50 | 28 |
| PIF4-mVenus | - | 2.85 | 0.11 | 79 |
| | + free mCherry | 2.64 | 0.06 | 34 |
| | + PLT3-mCherry | 2.47 | 0.11 | 44 |
| | + PLT3ΔPrD-mCherry | 1.53 | 0.32 | 24 |

**Supplementary table 17 SAXS data collection and analysis**

| Data-collection parameters | | |
|---|---|---|
| Instrument: | ESRF BM29 | |
| Wavelength (Å) | 0.99 | |
| q-range ($Å^{-1}$) | 0.007-0.5 | |
| Sample-to-detector distance (m) | 2.6 | |
| Concentration range (mg/mL) | 4 | |
| Temperature (K) | 277-300 | |
| Detector | Pilatus P3-2M | |
| Flux (photons/s) | $1*10^{13}$ | |
| Beam size (µm) | 500*200 | |
| SEC-SAXS | 1000 frames at 2 Seconds per frame | |
| **Structural parameters** | **ELF3 PrD** | **PLT3 PrD2** |
| I0 (kDa) [from Guinier] | 103 | 8.06 |
| Rg (Å) [from Guinier] | 80.1 | 34.3 |
| Volume ($Å^3$) | $2.1*10^6$ | $8.8*10^4$ |
| $D_{max}$ (Å) | 299 | 176 |
| Calculated MW from Vc (KDa) | 850 | 21 |
| Calculated theoretical MW monomer (kDa): | 29.15 | 22.47 |
| **software employed** | | |
| Primary data reduction: | BM29 autoprocessing pipeline | BM29 autoprocessing pipeline, FREESAS (Kieffer 2021) |
| Data processing | Scatter IV (Tully 2021) | |
